# A late cytoplasmic surveillance pathway ensures ribosome integrity

**DOI:** 10.1101/2025.10.29.685433

**Authors:** Ruta Chitale, Kaoling Guan, Shilpa Rao, Caroline Wang, Can Cenik, David W Taylor, Arlen W Johnson

## Abstract

Errors in ribosome assembly can produce defective subunits that can lead to aberrant translation events. How faulty ribosomes are recognized and whether they are recognized during assembly or translation remains poorly understood. We utilized a mutation in the ribosomal protein uL16 to track defective 60S subunits through the biogenesis and translation pathways. This mutation deletes a critical loop in the P site and arrests pre- 60S particles during late cytoplasmic maturation. However, simultaneous mutations in the late biogenesis factors Nmd3 and Tif6 bypass this block, releasing defective ribosomes into the translational pool. Cryo-EM and selective ribosome profiling reveal that these ribosomes can form peptide bonds, but stall predominantly at the first few codons. We show that the uL16 mutant ribosomes are detected and targeted for degradation during biogenesis and that they escape degradation if they enter translation. We identify Reh1 as a non-canonical ribosome assembly factor that is required for this surveillance pathway.

## Introduction

Ribosome biogenesis is a fundamental and highly orchestrated process that begins in the nucleolus with co-transcriptional rRNA processing and continues through a series of maturation events in the nucleus and cytoplasm^1,2^. These steps involve the hierarchical assembly of ribosomal proteins with rRNA, dynamic interactions with biogenesis factors, and extensive structural remodeling, culminating in the production of translationally competent 40S and 60S subunits^3^. Due to its complexity, ribosome biogenesis is highly sensitive to disruption. Defects in rRNA folding^4^, transcription^5^, post-transcriptional modifications^6,7^, translation^8^, and post-translational modifications^9^ can lead to the formation of aberrant intermediates. These defective ribosomes can compromise cellular homeostasis and have been implicated in a range of human diseases, including ribosomopathies and cancer^10–13^.

Cells have evolved robust quality control pathways to monitor translation^14^, including nonsense-mediated decay^15^, no-go decay^16^, and ribosome-associated quality control^17^, which recognize ribosomes stalled on faulty mRNAs and target them for degradation. These mechanisms act after ribosomes engage in translation. Despite compelling evidence of the existence of quality control pathways that detect nonfunctional ribosomes during assembly in the nucleus^18–21^, far less is known about how cells monitor the integrity of newly assembled subunits. Structural or functional defects in the ribosome itself are likely to cause stalling that is indistinguishable from mRNA-driven events^22^, raising the fundamental question of how cells distinguish between defective ribosomes and defective mRNAs.

In the cytoplasm, the last step of 60S assembly is the insertion of uL16 which completes the peptidyl transferase center (PTC)^23–25^ and triggers the release of the assembly factors Nmd3 and Tif6. Nmd3 is an essential nuclear export adaptor for the large subunit^26,27^ and Tif6 blocks recruitment of the 40S subunit^28–30^. The release of Tif6 by the GTPase Efl1 and its cofactor Sdo1 licenses the subunit for translation^31^. As Efl1 is a paralog of the translation elongation factor eEF2 and Sdo1 binds in the P-site, the activation of Efl1 by Sdo1 to release Tif6 has been proposed as a “test drive” which mimics aspects of itranslation elongation. We showed previously that the integrity of the P-site is assessed during the “test drive” and that defects in uL16 arrest biogenesis at this step^32^. However, the fate of these arrested subunits has not been characterized.

Most studies examining the fate of defective ribosomes have used mutations in ribosomal RNA. Mutations in 18S rRNA, including A1824C and A1755C which impair decoding, are recognized during translation through the 18S nonfunctional rRNA decay pathway ^19,20,33^. In contrast, mutations in the large subunit which impact the catalytic function of the ribosome, A2820G and U2954A in yeast, differentially affect progression of the subunits into translation. These 25S rRNA mutants are ultimately targeted for degradation by the proteasome^34,35^. However, the mechanism of detecting these subunits remains unknown and it is difficult to conceive how mutations in the catalytic center can be identified by the cellular machinery without engaging in translation.

Recent work in human cells has identified ZNF574 as a factor that detects defective ribosomes during assembly. Depletion of ZNF574 was shown to stabilize a uL16 mutant that is incorporated into the ribosome but blocks the release of eIF6^36^. Structurally, uL16 is positioned near the A- and P-sites of the ribosome and contains a conserved 10-amino-acid loop (M102-Q112) that is thought to contribute to proper tRNA positioning during peptide bond formation^37,38^. Prior work from our lab demonstrated that deletion of this loop is lethal, highlighting its essential role in ribosome function in yeast^39^. Although deletion of ZNF574 stabilizes mutant uL16-containing ribosomes, ZNF574 is not conserved throughout eukaryotes and how it interacts with nascent ribosomes to promote their degradation is unknown.

To investigate when and how defective assembled ribosomes are recognized, we have used the same uL16 mutant in yeast. We demonstrate that ribosomes lacking the uL16 P-site loop stall during late cytoplasmic biogenesis. We found that simultaneous mutations in *NMD3* and *TIF6* bypassed the assembly block, allowing the mutant ribosomes to progress into the translational pool. Using this genetic system, we asked whether defective subunits were subject to surveillance primarily during biogenesis or translation. We found that uL16 mutant ribosomes are detected and targeted for degradation during biogenesis before they enter the translation cycle. In addition, we previously identified Reh1 as the last assembly factor to be released from the pre-60S subunit^40^ and suggested that Reh1 could have a role in flagging defective subunits. Indeed, here we identify Reh1 as a key player in this surveillance pathway. These findings uncover a previously unappreciated layer of quality control during ribosome biogenesis and provide a model to dissect how cells recognize and eliminate defective ribosomes before and after translation initiation.

## Results

### Establishing a defective ribosome model

To investigate how cells handle defective ribosomes, we established a reporter system to examine quality control mechanisms during ribosome biogenesis and translation. We focused on the ribosomal protein uL16 (yeast Rpl10), which incorporates late during cytoplasmic 60S subunit maturation to complete the PTC and assembly of the subunit (**Fig. 1a**). This makes uL16 an ideal target because it avoids perturbing earlier assembly steps. Specifically, we used a mutant of uL16 lacking the P-site loop (residues 102–112), which interacts with P-site ligands (**Fig. 1b**). Deletion of the loop is lethal in yeast, highlighting its critical role in ribosome function^39^. To overcome lethality, we ectopically expressed a C-terminally Flag-tagged uL16 P-site loop mutant (uL16^mut^), which conferred a dominant-negative growth defect (**Fig. 1c**). We previously showed that this mutant arrests 60S maturation and blocks the recycling of Nmd3 and Tif6, however, the precise arrest point in the maturation pathway had not been determined. To characterize this mutant further, we analyzed the sedimentation pattern of the uL16^mut^ on sucrose density gradients. Unlike endogenous uL16, which distributes across the 60S, 80S, and polysome fractions, or Nmd3, which co-sediments exclusively with the 60S subunit, the uL16^mut^ accumulated predominantly in the 60S fraction (**Fig. 1d**), indicating a defect in the release of subunits into translation or a defect in subunit joining.

**Figure 1:**
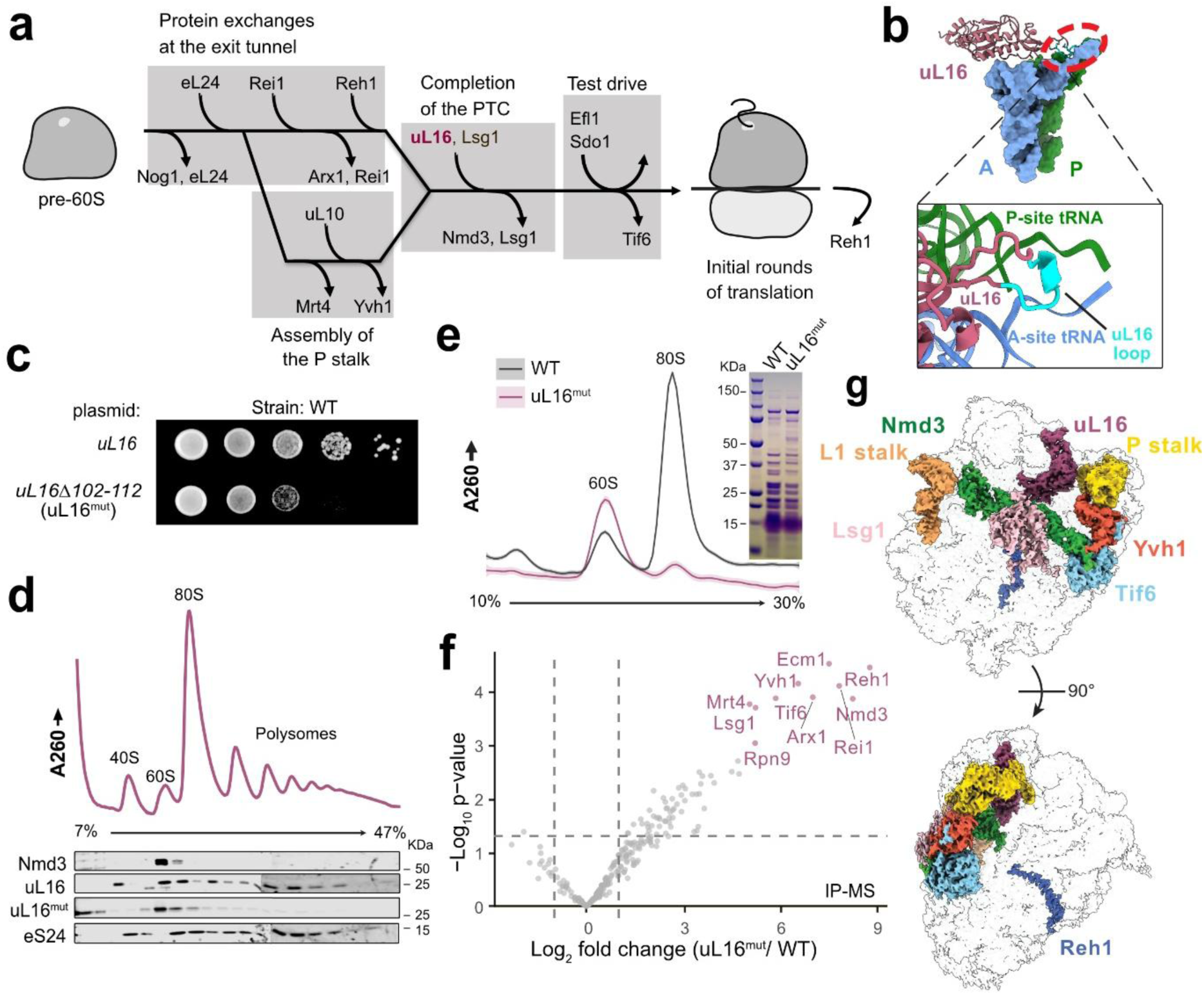
uL16^mut^ are trapped in the biogenesis pathway. **(a)** Simplified pathway of cytoplasmic maturation of the 60S ribosomal subunit in yeast, showing protein exchanges at the exit tunnel, assembly of the P stalk, completion of the PTC by uL16, test drive, and removal of the last biogenesis factor Reh1 during early rounds of translation. **(b)** P-site loop [amino acids 102-112] (cyan) of uL16 (mauve) interacts with A (blue) and P (green) site tRNAs (PDB:8UTI). **(c)** Serial dilutions of uL16 wildtype and uL16^mut^ expressed on a plasmid in wild type yeast (BY4741). **(d)** uL16^mut^ accumulates at 60S. Extract of wildtype cells (BY4741) expressing 3xFLAG-tagged uL16^mut^ was separated by sucrose density gradient sedimentation. UV trace monitoring A260 is shown. Fractions were analyzed by western blotting for the presence of uL16^mut^, uL16 (endogenous), eS24 and Nmd3. **(e)** uL16^mut^ predominantly associates with pre-60S particles. FLAG-tagged wild-type and uL16^mut^ were expressed in wild-type cells and affinity-purified. Left: SDS-PAGE showing pulldown of uL16. Right: A260 UV profiles of purified ribosomal complexes from wildtype vs. mutant samples sedimented through sucrose density gradients. **(f)** Volcano plot comparing results of LC-MS/MS from Flag-tagged uL16 wildtype and uL16^mut^ pull-down samples. p values (student’s t-test) and fold changes are shown. Proteins of interest are highlighted. **(g)** Cryo-EM map of affinity purified uL16^mut^ pre-60S ribosomes showing associated biogenesis factors, subunit joining surface (left), solvent-exposed surface (right).

To identify the specific maturation step disrupted by the uL16^mut^ and to identify associated biogenesis factors, we performed mass spectrometry on affinity-purified particles. Ribosomes containing Flag-tagged wild-type or uL16^mut^ were immunoprecipitated and separated on sucrose density gradients. Compared to wild-type, the mutant samples predominantly contained 60S particles and revealed a clear absence of 40S and depletion of 80S particles (**Fig. 1e**). Mass spectrometry of the free 60S fraction revealed a marked enrichment of the late 60S biogenesis factors Nmd3, Tif6, Lsg1, Yvh1, and Reh1 (**Fig. 1f**), indicating a defect in the final maturation stages of the 60S subunit, before the “test drive” step.

To investigate the structural consequences of uL16^mut^ deletion, we performed single-particle cryo-electron microscopy (cryo-EM) on ribosomes purified from cells expressing Flag-tagged uL16^mut^. We determined a 2.73 Å resolution structure of a pre-60S particle (**Supplementary Figure. 1, 3, supplementary Table 4**). Continuous density for the truncated uL16 loop was evident, indicating that the population of particles was relatively homogeneous. Consistent with our mass spectrometry results, this structure was enriched for the biogenesis factors Nmd3, Tif6, Lsg1, and Yvh1 at the subunit joining interface, with the assembly factor Reh1 in the exit tunnel (**Fig. 1g**). Yvh1 is a dual-specificity phosphatase known to promote P stalk assembly. The presence of Yvh1 in this structure was unexpected because our current understanding of 60S maturation is that the assembly of the P stalk is independent of assembly of the PTC or removal of Nmd3^23,24^. The accommodation of uL16 has previously been reported to be necessary for the release of Nmd3^23,32^, consistent with the retention of both Nmd3 and Tif6 on the pre-60S particle. Thus, uL16^mut^ blocks the removal of Nmd3, thereby impairing the "test drive" step that licenses the large ribosomal subunit for translation. This system enables us to trap mutant ribosomes within the biogenesis pathway, providing direct access to the quality-control mechanisms that operate during maturation.

### Mutations in Nmd3 and Tif6 relieve the biogenesis arrest of the uL16 mutant

Recurrent mutations in human uL16 have been identified as driver mutations in pediatric T-cell acute lymphoblastic leukemia^41^. These mutations are predominantly localized to arginine 98 (R98), with a single reported case involving a Q123P mutation. Notably, both residues lie at the base of the P-site loop of uL16. We previously demonstrated that these T-ALL-associated uL16 mutants impaired the release of Nmd3 from the nascent subunit and that this defect can be bypassed by mutations in Nmd3 that weaken its affinity for the ribosome and facilitate its release^42^. One such mutant is *nmd3-Y379D*, which partially disrupts the interaction between the eIF5A-like domain of Nmd3 and the ribosomal E site (**Fig. 2a, top inset**). Since the P-site loop of uL16 is also likely to impinge on the function of Sdo1 and Efl1 in the release of Tif6, we reasoned that this defect could be bypassed by mutations in Tif6 that are known to suppress loss of Sdo1. Here, we used the *tif6- V192F* mutation, which compromises Tif6 binding to uL14^43,44^ (**Fig. 2a, bottom inset**), allowing the mutant Tif6 to be released independently of Sdo1 and Efl1, while retaining Tif6 function in assembly. We tested whether bypass mutations in *NMD3* and *TIF6* could suppress the dominant negative effect of uL16^mut^. We found that *nmd3-Y379D* and *tif6- V192F* individually partially suppressed the growth defect of the loop deletion mutant, with the combination of both mutations producing robust rescue (**Fig. 2b**).

**Figure 2:**
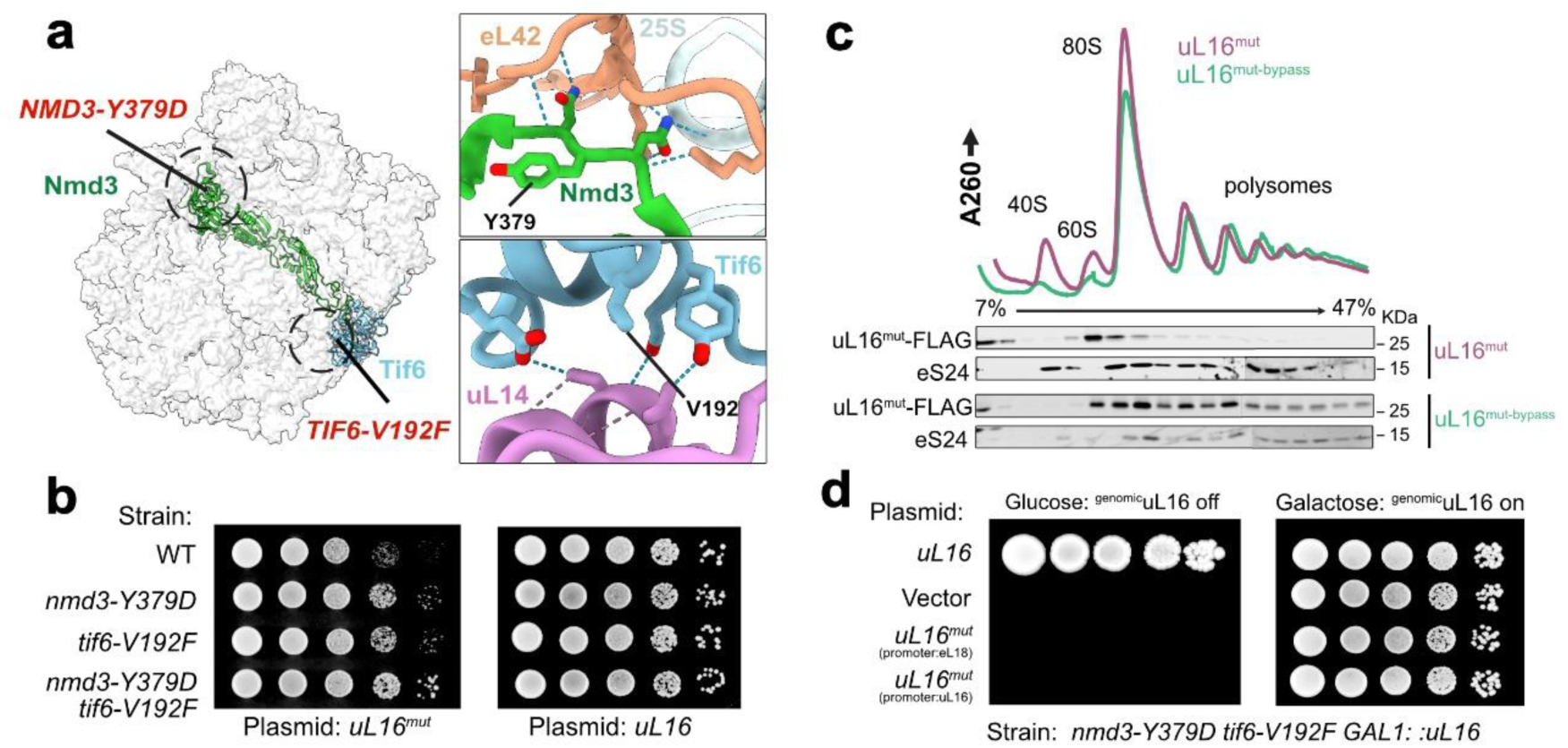
Mutations in *NMD3* and *TIF6* release uL16^mut^ into translation. **(a)** Structure of the subunit joining surface of a pre-60S showing Nmd3 and Tif6 with positions of the suppressor mutations *nmd3-Y379D* and *tif6-V192f* (left). Inset, top: Y379D in Nmd3 disrupts interactions with eL24and 25S rRNA, indicating its role at the Nmd3-60S interface. Inset, bottom: V192F in Tif6 disrupts interaction with uL14 (PDB:6N8O). **(b)** Serial dilutions of single and double mutants of *NMD3* and *TIF6* expressed with plasmid-borne uL16 wildtype and uL16^mut^ demonstrate suppression of slow growth phenotype of uL16^mut^. **(c)** Sucrose density gradient sedimentation of Wildtype and *nmd3-Y379D tif6- V192F* cells when expressed with plasmid-borne uL16^mut^ reveals sedimentation into polysome fractions. UV trace monitoring A260 is shown. Fractions were analyzed by western blotting for the presence of uL16^mut^ FLAG, and eS24. **(d)** Serial dilutions showing yeast strain *nmd3-Y379D tif6-V192f* with genomic *GAL1:uL16*, expressing *uL16*, empty vector or *uL16^mut^* under the control of the *eL18* or *uL16* promoters, and growing on glucose (genomic *GAL1:uL16* repressed) or galactose (expressed) media.

Given that the *nmd3-Y379D* and *tif6-V192F* mutations bypassed the dominant negative phenotype caused by arresting 60S maturation, we considered that these mutations may allow the mutant ribosomes to escape the licensing step and enter translation. To test this, we examined the sedimentation of uL16^mut^ in the *nmd3-Y379D tif6-V192F* mutant strain (hereafter referred to as the bypass strain) using sucrose density gradient centrifugation. While a fraction of uL16^mut^ ribosomes still sedimented in the 60S fraction, the majority of the mutant protein was observed in the 80S and polysome fractions in the bypass background (**Fig. 2c**), indicating escape from the maturation arrest and engagement with 40S subunits. Similar to the partial rescue of the dominant negative effect on growth, we observed that the individual suppressing mutations each showed partial release of defective ribosomes from the biogenesis arrest (**Supplementary Figure. 5a, b**). These findings suggest that Nmd3 and Tif6 act as gatekeepers that prevent the release of defective pre-60S subunits. The dominant-negative effect of uL16^mut^ is likely due the failure to recycle the key biogenesis factors Nmd3 and Tif6, an effect that is alleviated in the bypass strain. Complementation analysis revealed that uL16^mut^ was unable to rescue the lethal phenotype in the bypass strain, indicating that although the mutant ribosomes were engaging in translation, they were unable to support growth (**Fig. 2d**).

### Bypass mutations restore late 60S maturation

To investigate the nature of the 60S particles that escape maturation, we performed mass spectrometry of ribosomes from the 60S and 80S fractions. Ribosomes were affinity-purified from the bypass strain and separated by sucrose gradient centrifugation to isolate the 60S and 80S fractions (**Supplementary Figure. 6a**). Analysis of the 60S fraction revealed enrichment of several 60S biogenesis factors, with particularly high enrichment of Reh1 and Tif6 (**Supplementary Figure. 6b**). This pattern differed from that observed in the 60S fraction of the wild-type background expressing uL16^mut^, where subunits were enriched for a broader range of late-stage maturation factors. In contrast, the 80S fraction from the bypass strain exhibited a near-complete absence of biogenesis factors, with the exception of Reh1 (**Supplementary Figure. 6c**). These findings suggest that suppression by *nmd3-Y379D* and *tif6-V192F* enables bypass of the biogenesis defect by promoting the release of Nmd3 and Tif6.

To determine whether the bypassed uL16^mut^ ribosomes are translationally competent, we performed single particle cryo-EM on ribosomes isolated from the bypass background. This analysis revealed that more than half of the particles corresponded to free 60S subunits (∼203,066 particles), while the remaining ∼187,670 particles represented assembled 80S ribosomes (**Supplementary Figure. 2, supplementary Table 4**). Focused 3D classification on Nmd3 on the free 60S subunits yielded a 2.83 Å resolution structure of a pre-60S particle containing the biogenesis factors Nmd3, Lsg1, Tif6, and Reh1 (**Supplementary Figure. 6d**). In addition, we resolved a 2.78 Å structure of a distinct pre-60S particle that lacked Lsg1 and Nmd3 but retained Tif6 at high occupancy, along with Reh1, whose C-terminus remained inserted into the exit tunnel (**Supplementary Figure. 6e**). This structure likely represents a particle that has progressed beyond Nmd3 release and is poised for Sdo1 binding, a prerequisite for Tif6 removal and subsequent subunit joining. In the bypassed structure containing biogenesis factors, the C-terminus of Nmd3 (residues 409–500) is visible (**Supplementary Fig. 6d**). This region, which has not been resolved in previous structures, folds into a small domain that replaces helix 38 (H38) of the 25S rRNA (**Supplementary Fig. 6f**). The presence of the Nmd3 C-terminus coincides with the absence of Yvh1, suggesting an intermediate state that occurs after Yvh1 removal.

We identified a well-defined density corresponding to the C terminus of Reh1 extending into the polypeptide exit tunnel. Additional density on the Tif6 surface was consistent with the N-terminal region of Reh1, and AlphaFold3 modeling assigned the first zinc cluster to this region (**Supplementary Fig. 7a-c**). The Reh1 density extends from Tif6 to contact eL24, traverses adjacent rRNA elements, and forms additional interactions via its second and third zinc clusters with eL22, which could also be modeled using AlphaFold3 (**Supplementary Fig. 7b, d**). The density then continues along eL19 and uL23 before entering the exit tunnel (**Supplementary Fig. 7e**). The trajectory of Reh1 closely parallels that of its paralog, Rei1, except that residues 388–377 of Reh1 approach H7 and uL29, whereas Rei1 diverges away from H7^45^; both, however, follow a shared path across eL22. Together, these results indicate that bypass mutations relieve the late 60S maturation block by promoting the sequential release of Nmd3 and Tif6, thereby enabling productive subunit joining and formation of translationally competent 80S ribosomes.

### uL16 mutants in the bypass strain actively engage in translation

Notably, approximately 50% of the total particles isolated from the bypass background corresponded to mature 80S ribosomes, supporting a model in which a substantial fraction of bypassed mutant ribosomes advance beyond the biogenesis defect and enter the translational pool. We analyzed the 80S ribosomal particles from the bypass strain to assess their translational state. Using a focused 3D classification strategy based on tRNA occupancy, we identified four distinct ribosomal complexes (**Supplementary Fig. 2a, 4, supplementary Table 4**). The first structure, resolved at 2.66 Å, contained mRNA and tRNAs in the classical A/A and P/P positions, with an empty E site and an open L1 stalk (**Fig. 3a, b**). The second structure, resolved at 2.82 Å, contained a single tRNA in the P/P-site, along with mRNA, with both the A and E sites unoccupied (**Fig. 3d, e**). The third structure, resolved at 3.02 Å, represented a pre-translocation intermediate, with tRNAs in A/P and P/E hybrid positions and L1 stalk engaging with the P/E tRNA (**Fig. 3g, h**). Although cells were treated with cycloheximide, which typically traps ribosomes in a non-rotated state, this complex was found in a rotated hybrid state, potentially reflecting slowed tRNA translocation due to the absence of the uL16 P-site loop. The final structure was an empty 80S ribosome, lacking both tRNAs and mRNA. Together, these structures demonstrate that in the bypass strain, 80S ribosomes are capable of engaging in translation despite the absence of the P-site loop of uL16.

**Figure 3:**
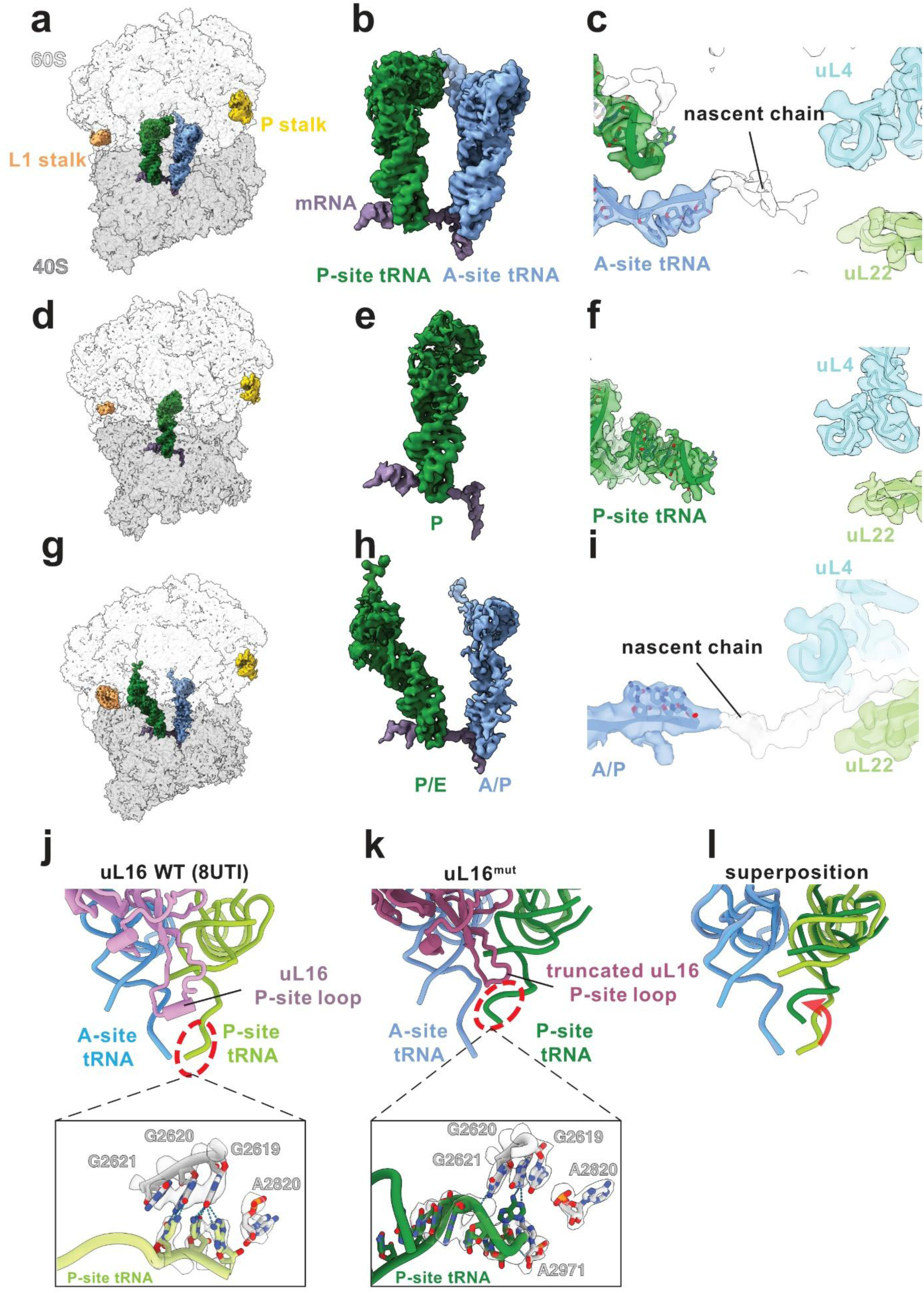
Cryo-EM structures of translating ribosomes containing uL16^mut^. Affinity- purified uL16^mut^ mutant ribosomes from the bypass strain were imaged on a Titan Krios (FEI) and analyzed with CryoSPARC, revealing four distinct 80S structures. **(a)** Cryo-EM model and map of an 80S ribosome with both A-site and P-site tRNAs in classical A/A and P/P conformations. The L1 stalk and P stalk are indicated. **(b)** Corresponding tRNA and mRNA density for the ribosome in (a), highlighting the conformation of A/A and P/P tRNAs and positioning of the mRNA. **(c)** Density for the nascent polypeptide chain extending from the A/A tRNA. **(d)** 80S ribosome with only a classical P-site tRNA (P/P conformation). **(e)** Corresponding tRNA and mRNA density map for the ribosome in (d). **(f)** Density for nascent chain associated with the P/P tRNA. **(g)** 80S ribosome with hybrid- state A/P and P/E site tRNAs. **(h)** Corresponding tRNA and mRNA density maps showing hybrid tRNA conformations. **(i)** Density for the nascent chain extending from the A/P tRNA. In all panels, the ribosomal exit tunnel is highlighted by uL4 and uL22, which line the wall of the tunnel. **(j)** In wild-type ribosomes, the P-site loop of uL16 contacts the acceptor stem of the P-site tRNA. Inset: In wild-type ribosomes, a classical hydrogen bond network is observed between the P-site tRNA and the P-loop of the 25S rRNA. **(k)** In the absence of the uL16 P-site loop, the acceptor stem of the P-site tRNA retracts and loses interaction with the P-loop of the 25S rRNA in the structures containing either A/A and P/P tRNAs or P/P tRNA alone. Inset: In the absence of the uL16 P-site loop, the P-site tRNA adopts novel interactions with surrounding rRNA, presenting suboptimal positioning of the P-site tRNA for peptidyl transfer. **(l)** Superposition of A- and P-site tRNAs from wild-type and uL16^mut^ ribosomes shows altered positioning of the P-site tRNA in the mutant.

To determine whether these ribosomes are competent for polypeptide synthesis, we assessed the presence of densities corresponding to nascent peptide across the different structural states. In the classical state, which contains A/A and P/P tRNAs, we observed a small but distinct density extending from the 3′ end of the A-site tRNA, corresponding to the nascent polypeptide chain (**Fig. 3c**). Although the exact length of the nascent chain is difficult to determine, the density appears to represent more than two amino acids, suggesting that the ribosome is in a non-rotated, post-peptide bond formation state of translation. The ribosomal state containing only a P-site tRNA exhibits a small density at the 3′ end of the tRNA and an empty exit tunnel, which may correspond to initiator tRNA (**Fig. 3f**). In contrast, the hybrid-state ribosomes, which contain A/P and P/E-site tRNAs, display an extended density from the 3′ end of the A/P tRNA and projecting into the ribosomal exit tunnel (**Fig. 3i**). As the distance between the tRNA and the entrance of the exit tunnel corresponds to approximately six amino acids, the observed density reflects a nascent chain of at least this length. However, interpretation of the density in the exit tunnel is restricted by the presence of Reh1 in the exit tunnel in approximately half of the particles, making it difficult to unambiguously assign the tunnel density to either Reh1 or the nascent peptide. Nevertheless, the presence of a polypeptide chain extending beyond the PTC indicates that peptide bond formation has occurred, supporting the conclusion that these mutant ribosomes retain the ability to catalyze peptide bond formation. However, the rate and overall efficiency of translation is likely to be severely compromised.

A previous study from our lab showed that Reh1 uniquely associates with nascent 80S subunits as they enter the translation cycle^40^. To independently validate this observation and confirm that the ribosomes we observe originate from the biogenesis pathway rather than recycled subunits, we reexamined all 80S particles for presence of Reh1 (**Supplementary Figure. 2b (i)**). Approximately half of the particles exhibited alpha-helical density within the exit tunnel. However, because many of these ribosomes are actively translating and contain nascent polypeptides, unambiguously assigning this density to Reh1 is challenging. To refine our analysis, we further classified Reh1 containing particles based on the presence of tRNAs (**Supplementary Figure. 2b (ii), supplementary Table 4**). About 37% of the Reh1 bound particles lacked tRNAs (“empty” ribosomes), allowing us to tentatively attribute the exit-tunnel density to Reh1 in these particles (**Supplementary Fig. 8b**). In contrast, the remaining particles contained tRNAs. Among these, we focused on ribosomes bearing only a P-site tRNA with a short nascent chain that did not appear to reach the exit tunnel. In these particles, the alpha-helical density within the exit tunnel is again likely to be Reh1 (**Supplementary Fig. 8c**). However, in both cases, the density lacked the well-defined features corresponding to bulky side chains seen for Reh1 in the 8UTI structure, suggesting that if this density represents Reh1, it may adopt alternative conformations within the exit tunnel (**Supplementary Fig. 8a-c**).

Lastly, we observed the N-terminus of eS25 intercalating between the anticodon stem-loop helices of the A- and P-site tRNAs. This interaction was previously reported by our lab in Reh1-containing 80S ribosomes, where we speculated that eS25 may play a role in early elongation^40^. A systematic analysis of structures with different tRNA configurations revealed that the N-terminus of eS25 is absent in initiating ribosomes containing only a P-site tRNA, as well as in ribosomes in the hybrid A/P and P/E state. In contrast, it is consistently present in ribosomes with classical A- and P-site tRNAs. Taken together, these results reveal that the uL16^mut^ ribosomes in the bypass state are actively engaging in translation and are competent for polypeptide synthesis.

### The P-site loop of uL16 positions the P-site tRNA

The P-site loop of uL16 forms specific contacts with the acceptor stem of the P-site tRNA and is thought to facilitate its proper positioning within the PTC^37,38^. Deletion of this loop is lethal, however, using our genetic system, we were able to bypass this biogenesis arrest and capture ribosomes lacking the P-site loop during translation, thereby enabling structural dissection of the resulting defects. Our initial analysis of 80S ribosome structures revealed no obvious global structural abnormalities. To more directly assess the functional role of the P-site loop within the catalytic center, we next examined the architecture of the PTC in the mutant ribosomes and its impact on tRNA positioning. In wild-type ribosomes, the P-site loop of uL16 makes stabilizing contacts with the P-site tRNA, including a hydrogen bond between residue R110 and the tRNA’s acceptor stem (**Fig. 3j**). In contrast, deletion of the P-site loop abolishes these interactions, resulting in a distorted conformation and retraction of the acceptor stem of the P-site tRNA (**Fig. 3k, l**). Proper positioning of the P-site tRNA within the PTC is further supported by several rRNA elements. In wild-type ribosomes, the P-site tRNA interacts with the P-loop (A2620, A2619) and with the universally conserved nucleotide in the catalytic center, A2820 (A2451 in *E. coli*) (**Fig. 3j, inset**). However, in the absence of the P-site loop of uL16, the retracted tRNA conformation disrupts these canonical contacts (**Fig. 3k, inset**). Instead, the tRNA establishes new interactions with A2621, also part of the P-loop, and with A2971, which appears to stabilize this alternate, retracted configuration. We speculate that mispositioning of the P-site tRNA impairs peptide bond formation and reduces elongation efficiency, ultimately contributing to the lethal phenotype observed in bypassed mutant cells.

### Footprint analysis reveals limited translation by the loop mutants

To better understand how P-site tRNA mispositioning contributes to translational defects, we performed selective ribosome profiling on bypassed mutant ribosomes lacking the uL16 P-site loop. Cell extracts were prepared under the same conditions used for cryo-EM to correlate changes in ribosome footprints with structural changes. Ribosome profiling libraries were prepared before purification (Total) and after affinity-purification of FLAG-tagged uL16^mut^ mutant ribosomes (IP) (**Fig. 4a**). In both datasets, the vast majority of reads (∼99% in Total and ∼94% in IP) mapped to coding sequences (CDS), indicating efficient capture of translating ribosomes (**Fig. 4b**). Moreover, ribosome footprint counts per gene were highly correlated between biological replicates (Spearman’s rank correlation, ρ = 0.971 for Total and 0.950 for IP), suggesting reproducibility of the ribosome profiling data.

**Figure 4:**
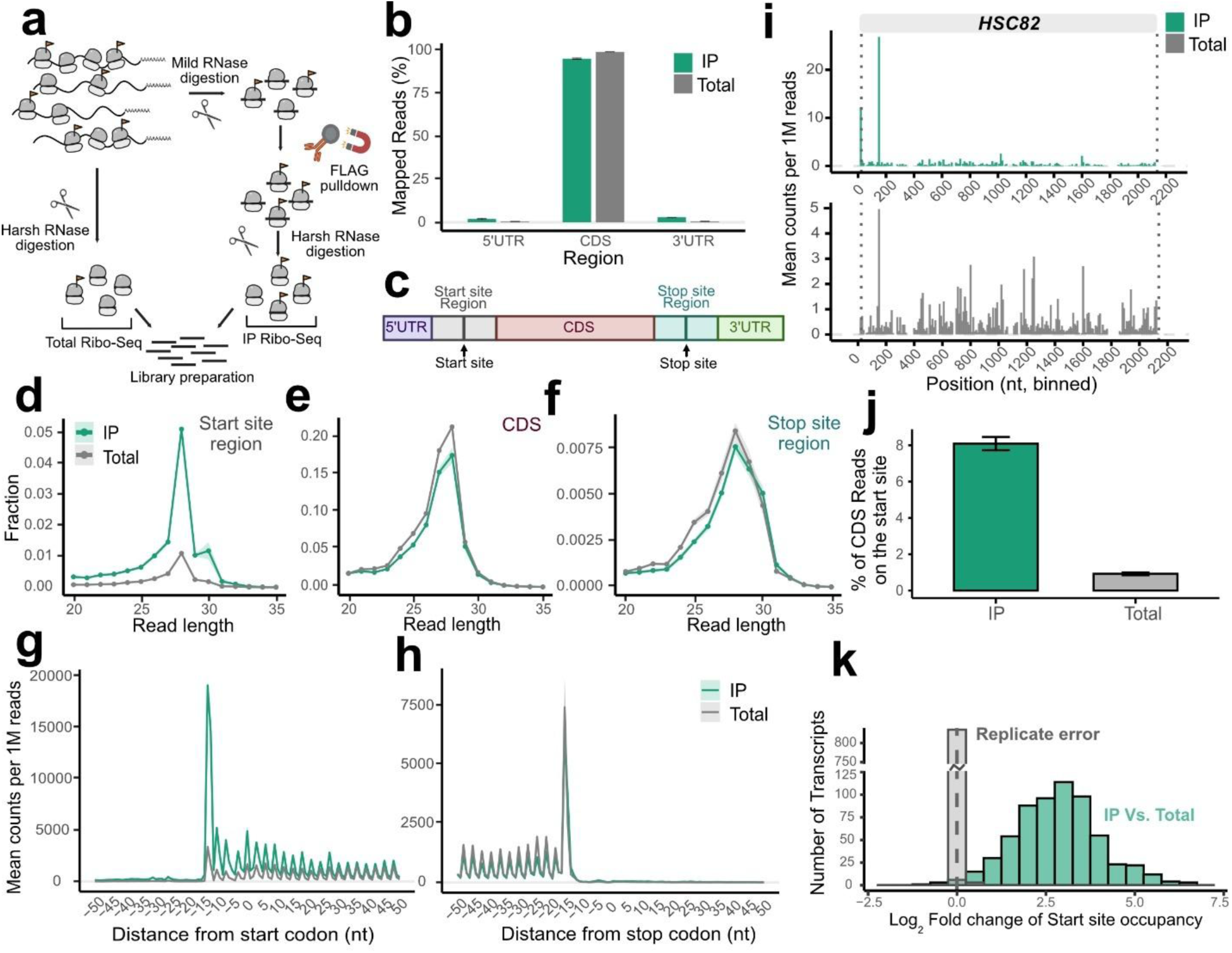
uL16^mut^ ribosomes are enriched at the start codon. **(a)** Schematic of the selective ribosome profiling workflow. **(b)** Bar graph showing the percentage of ribosome profiling reads mapping to the 5’ UTR, CDS, and 3’ UTR for both Total and IP samples across biological replicates (n = 3). **(c)** Schematic representation of transcript regions and junction definitions used for metagene analyses. **(d–f)** Distribution of read lengths in selective (IP) and total ribosome profiling libraries from the uL16^mut^, plotted separately for reads mapping to the start site region (d), CDS (e), and stop site region (f). Values represent means ± SE from three replicates. **(g)** Metagene analysis of ribosome footprints around start codons for Total (grey) and IP (green) samples. Read density (counts per million) is plotted as a function of nucleotide position for read length 26-29nt. **(h)** Metagene analysis of footprints around stop codons. **(i)** Distribution of 28-nt RPFs on the HSC82 transcript from purified uL16^mut^ IP and Total ribosome libraries. Data are P-site adjusted. Dotted lines indicate annotated translation start and stop sites. **(j)** Percentage of CDS-mapping reads that align precisely with the start codon, shown for Total and IP samples. **(k)** Histogram of changes in start site occupancy between IP (purified mutant ribosomes) and Total ribosomes. Data represents average start ratios from biological replicates.

To assess translation states, we examined the length distribution of ribosome protected fragments across different transcript regions (**Fig. 4c**). Within CDS and near the translation stop site, we observed 28-nt footprints, consistent with classical ribosome profiles obtained from cycloheximide-treated samples containing an occupied A-site (**Fig. 4e, f**). For the region surrounding the translation start site, both Total and IP samples showed the expected 28-nt peak; however, the IP sample also displayed an additional distinct 30-nt peak, suggesting altered ribosomal dynamics specific to the mutant ribosomes (**Fig. 4d, Supplementary Figure. 9a, b**). We speculate that this 30-nt footprint reflects the hybrid A/P P/E ribosomes that we observed by cryo-EM.

We next analyzed metagene coverage around the start and stop codons to assess global translation dynamics. Both Total and IP ribosome footprint samples displayed clear three-nucleotide periodicity, consistent with active translation (**Fig. 4g, h**). However, the IP sample showed pronounced accumulation of ribosome footprints at the start codon and within the first ∼15 codons (∼45 nucleotides) of the CDS, regardless of transcript length, indicating ribosome pausing or slowed early elongation (**Fig. 4g, i, Supplementary Figure. 9c**). In contrast, the Total sample showed a more uniform distribution of reads. Approximately 8% of IP reads mapped to the start codon, compared to only 0.86% in the Total sample (**Fig. 4j**). Similarly, 19.3% of IP reads were enriched within the first 15 codons, whereas only 3.5% of Total reads were found in this region, consistent with a defect in early translation.

We next assessed whether the observed metagene enrichment around the start codon is driven by a small number of genes or is a general feature of all. Start site occupancy was calculated as the ratio of reads at the start codon to total reads across CDS. Compared to the Total sample, the IP sample exhibited transcriptome-wide stalling at the start codon, as shown by the histogram of the start site occupancy (**Fig. 4k**). These observations suggest that uL16^mut^ ribosomes are impaired during the early steps of translation, particularly in initiation or forming the first few cycles of elongation. Although the mutant ribosomes were stalled primarily within the first 15 codons, we noted a substantial number of IP reads distributed across the CDS and the presence of ribosome footprints near stop codons (**Fig. 4h, Supplementary Fig. 9c**). These reads could reflect continued translation by mutant ribosomes once they have passed some threshold of approximately 15 codons. Alternatively, and perhaps more likely, is that these reads represent a low level of contamination from wild-type ribosomes. This interpretation is supported by our structural analysis. We found that approximately 89% of the ribosomes engaged with mRNA contain a short nascent chain that doesn’t reach the exit tunnel. Consequently, not more than 11% of the reads from the mutant ribosomes can be derived from ribosomes in downstream CDS. Thus, the impairment in early translation appears to be sufficient to disrupt overall protein synthesis and is likely insufficient to support normal cell growth.

### uL16 mutants are rapidly degraded in the biogenesis pathway

Having identified systems in which we can stall defective ribosomes during biogenesis or early in translation, we next asked if quality control mechanisms for defective ribosomes act primarily during biogenesis or translation. To this end, we monitored levels of the uL16^mut^ protein in both the biogenesis-arrested and translation-bypass states. In the biogenesis-arrested condition, mutant protein levels were reduced compared to wild-type, whereas levels were increased in the bypass strain, suggesting stabilization once the defective ribosomes escape biogenesis and enter translation (**Fig. 5a, b**). To further pinpoint when degradation occurs, we measured the half-lives of uL16^mut^ in both states. We found that the mutant protein is rapidly degraded in the biogenesis-arrested state but is relatively stabilized in the bypass strain, indicating that degradation is primarily triggered during ribosome biogenesis and that they are more stable once they enter translation (**Fig. 5c-e**). These observations suggest the existence of a dedicated degradation pathway operating during ribosome assembly.

**Figure 5:**
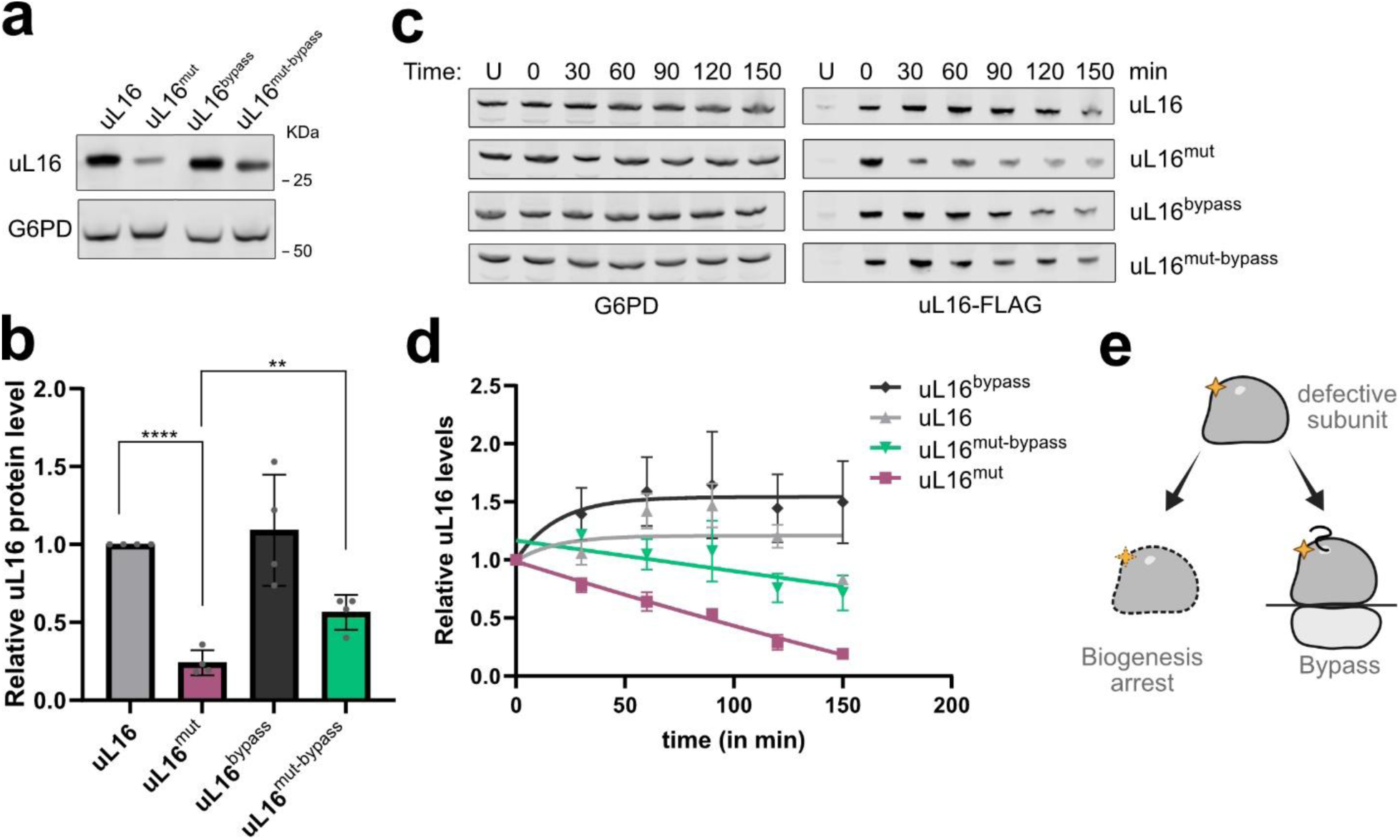
uL16^mut^ are degraded during ribosome biogenesis. **(a)** Western blot analysis of steady-state levels of FLAG-tagged uL16 wild-type (WT) and uL16^mut^ expressed in either WT or bypass (*nmd3Y379D tif6V192F*) strain backgrounds. G6PD serves as a loading control. **(b)** Quantification of uL16 protein levels from (a), normalized to G6PD and expressed relative to uL16 WT in WT cells. **(c)** Time course of uL16 degradation. FLAG-tagged uL16 WT or loop mutant was transiently expressed under the GAL1 promoter in WT or bypass strains. Transcription was repressed by the addition of glucose, and protein levels were monitored over time. **(d)** Quantification of uL16 degradation over time shown in (c). Protein levels were normalized to G6PD and plotted relative to levels at time zero (just after glucose addition). **(e)** Schematic illustrating the fate of ribosomal particles. Subunits that engage in the translation pathway are stabilized (as in the bypass condition), whereas particles that remain in biogenesis are directed for degradation.

### Reh1 uniquely marks defective nascent subunits for degradation

Since defective ribosomal subunits are degraded during biogenesis, we next asked which factors might contribute to tagging these subunits for quality control. To address this, we examined mass spectrometry data from the uL16^mut^ strain arrested during 60S biogenesis. This analysis revealed several associated biogenesis factors, including Nmd3, Tif6, Lsg1, Yvh1, and Reh1. Among these, Reh1 emerged as particularly noteworthy for several reasons. First, Reh1 is a late-acting biogenesis factor that remains bound to pre-60S subunits until the onset of translation, making it the last factor to dissociate^40^. Second, it localizes within the polypeptide exit tunnel, a site frequently targeted by ribosome quality control factors. We had previously speculated that the persistence of Reh1 could serve as a marker for identifying defective subunits destined for degradation. We therefore hypothesized that Reh1 may function in coupling late-stage biogenesis to quality control mechanisms.

To test for a functional interaction between Reh1 in uL16^mut^, we analyzed the effect of uL16^mut^ on cell growth of wild-type and *reh1Δ* mutant cells. Deletion of *REH1* markedly sensitized cells to expression of uL16^mut^, resulting in slower growth compared with uL16^mut^ in wild-type cells, indicating a genetic interaction (**Fig. 6a**). Strikingly, steady-state analysis revealed a ∼2.5-fold accumulation of uL16^mut^ protein in the *reh1Δ* background compared to wild-type (**Fig. 6b, c**).

**Figure 6.**
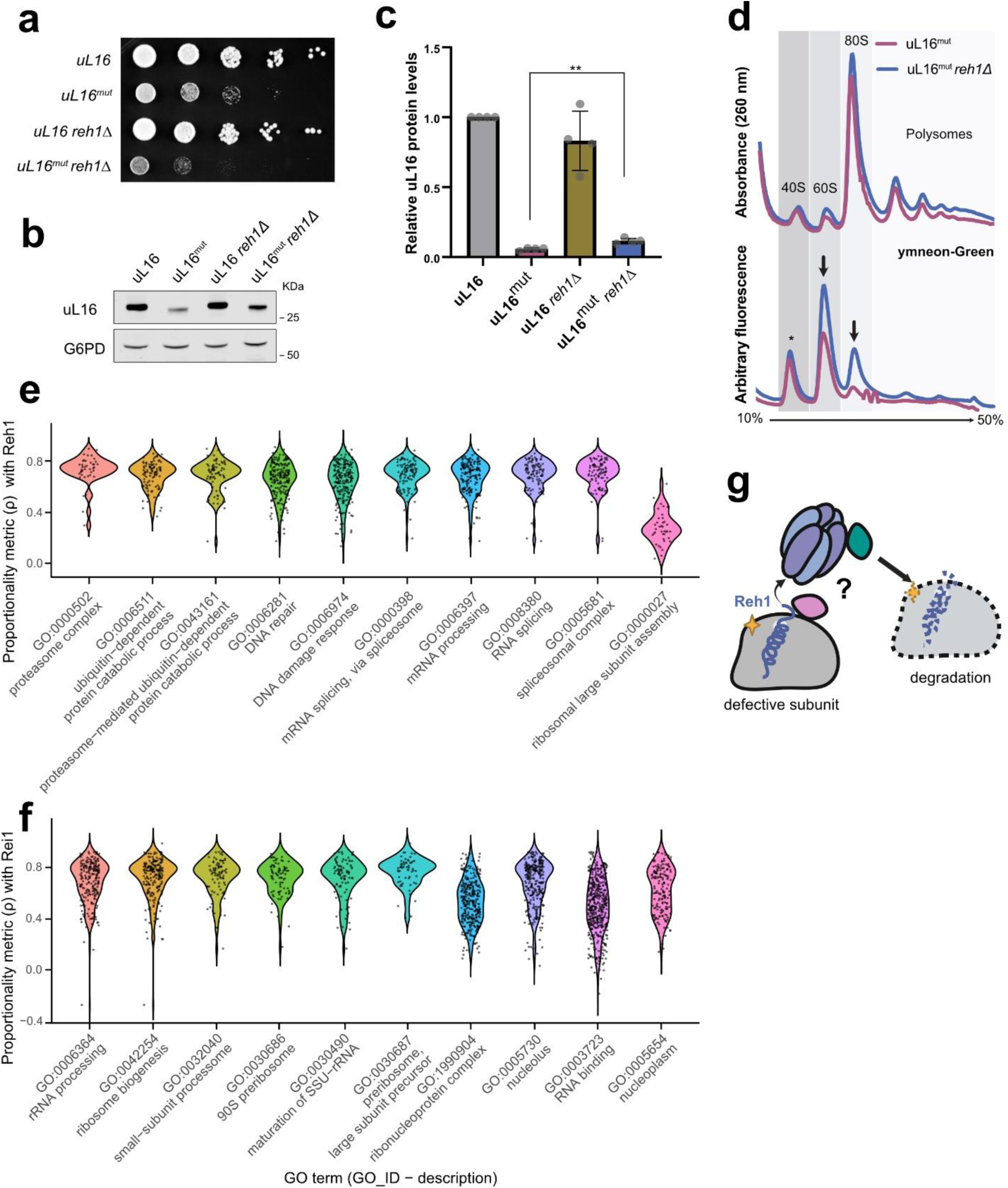
Biogenesis factor Reh1 stabilizes uL16^mut^. **(a)** Serial dilution growth assay of uL16 and uL16^mut^ expressed from a plasmid in the presence or absence of Reh1. **(b)** Western blot analysis of steady-state levels of FLAG-tagged uL16 wild-type (WT) and uL16^mut^ expressed in WT or *reh1Δ* strain backgrounds. G6PD serves as a loading control. **(c**) Quantification of uL16 protein levels from (b), normalized to G6PD and expressed relative to uL16^mut^ in WT cells. **(d)** uL16^mut^ predominantly accumulates in the 60S fraction in *reh1Δ* cells. Extracts from WT (BY4741) and *reh1Δ* strains expressing ymNeonGreen- tagged uL16^mut^ were separated by sucrose density gradient sedimentation. UV absorbance at 260 nm (top) and fluorescence corresponding to uL16 (bottom) are shown. * Indicates nonspecific fluorescence that sediments with 40S peak. **(e, f)** Co-expression analysis of Reh1 (e) and Rei1 (f) using gene-set enrichment analysis; the top ten most significantly enriched GO terms are shown. **(g)** Model for Reh1-mediated quality control of defective ribosomal subunits. Reh1 acts as a surveillance factor that flags defective or arrested subunits during late stages of 60S maturation, marking them for downstream quality control and degradation pathways.

To define the stage at which *reh1Δ* stabilizes uL16^mut^, we performed polysome profiling using a ymNeonGreen-tagged uL16^mut^ construct in the presence or absence of Reh1. Fluorescence and A260 profiles revealed that in reh1Δ mutant cells uL16^mut^ ribosomes accumulate predominantly in the 60S fraction, accompanied by a modest increase in 80S and polysome peaks (**Fig. 6d**). Because Reh1 is dispensable for ribosome biogenesis when Rei1 is present^46^, the increased 60S signal is unlikely to reflect a defect in subunit assembly. Instead, these data indicate that, in the absence of Reh1, uL16^mut^-containing pre-60S particles are stabilized. We speculate that the increased signal of uL16^mut^ in the 80S and polysome fractions in the *reh1Δ* mutant results from a low rate of Tif6 release due to the loss of Tif6 stabilization by Reh1. Notably, this profile differs from that of the bypass strain, where 80S and polysome levels increase while 60S levels decline, opposite to the pattern observed in *reh1Δ* cells. We speculate this difference reflects the efficiency of Tif6 release in the *reh1Δ* mutant compared to the bypass strain, in which Tif6 release is highly efficient. Together, these results indicate that Reh1 promotes turnover of uL16^mut^-containing pre-60S subunits, rather than acting solely on the uL16 protein itself.

Given that Reh1 functions in late-stage ribosome assembly, the stabilization observed for the uL16^mut^ suggested that Reh1 has adopted a quality control function and thus, may be regulated differently from canonical ribosome assembly factors. Because co-expression can indicate functional similarity, with genes participating in the same pathway or complex tending to be expressed under similar conditions, we examined whether Reh1 displays coordinated expression patterns with other genes. To investigate this possibility, we analyzed the co-expression profile of Reh1 to identify genes with coordinated expression patterns, using a yeast co-expression dataset from Rich et al.^47^. In this dataset, co-expression between ORFs was quantified using the proportionality metric (ρ) on clr-transformed RNA-seq data, which has been suggested to accurately capture functional similarities^48,49^. Using this dataset performed an unbiased gene set enrichment analysis to determine the biological processes most strongly associated with Reh1 co-expression. Contrary to expectation, Reh1 showed strong co-expression with genes involved in proteasome function, ubiquitin-dependent catabolic processes, mRNA splicing, and DNA repair, but exhibited only weak correlation with large ribosomal subunit assembly genes (**Fig. 6e**). To assess whether this pattern was unique to Reh1, we compared its co-expression profile with that of Rei1, a paralog of Reh1 that functions upstream of Reh1 in pre-60S assembly^50^. Rei1 displayed high co-expression with canonical ribosome biogenesis and rRNA processing genes, consistent with its established role in ribosome maturation (**Fig. 6f**). Together, these results suggest that despite their structural similarity, Reh1 may function in distinct or regulatory aspects of ribosome biogenesis, potentially linking late-stage ribosome assembly with protein quality control.

To determine whether Reh1 expression differs from that of other ribosome biogenesis factors, we performed a comparative co-expression analysis across hand selected gene ontology categories associated with ribosome biogenesis, translation, ubiquitination and proteasome. We compared Reh1 expression pattern with biogenesis factors acting at distinct stages of ribosome assembly, as well as with quality control and proteasomal components. This analysis allowed us to assess whether Reh1 clusters with canonical assembly factors or exhibits a transcriptional signature more consistent with quality control machinery. Notably, the co-expression pattern of Reh1 more closely resembled that of quality control and proteasomal factors, rather than canonical biogenesis factors, suggesting a potential role of Reh1 in quality control (**Supplementary Figure. 10**). Together, these findings suggest that Reh1 is more than just an assembly factor and functions at the interface between ribosome biogenesis and protein quality control, acting as a late-stage surveillance factor that helps identify and target defective subunits for degradation (**Fig. 6g**).

## Discussion

Here, we developed a system to test if quality control of nascent 60S subunits occurs during the licensing step of 60S maturation or during the initial rounds of translation elongation. Using a mutation in uL16 that impacts ligand binding in the P site, we found that the defective ribosomes are actively turned over when they stall at the licensing step. When these subunits are allowed to bypass this step and enter translation, they are relatively stabilized, despite the fact that they are still severely impaired for translation. Our results suggest that cells have evolved mechanisms to detect faulty 60S subunits, at least for the 60S mutant used here, prior to their engagement in translation. It is surprising that once engaged in translation these subunits become more stable because it is only during translation that the specific function of polypeptide synthesis can be assessed.

Additionally, we found that the turnover of defective subunits is dependent on Reh1. We previously showed that in yeast, Reh1 is the last ribosome assembly factor to be released from the newly made large subunit and that it is released after subunit joining, during translation^40^. In that work we had also provided evidence that the growing nascent chain evicts Reh1 from the exit tunnel. Thus, Reh1 seemed poised to uniquely identify the nascent subunit in its first round of translation and we speculated that its persistence on ribosomes could be a flag for identifying a defective subunit for subsequent degradation. In our current study, we find that deletion of *REH1* stabilizes defective nascent 60S subunits, allowing us to conclude that Reh1 is indeed a quality control factor for nascent 60S subunits. However, contrary to our expectations, Reh1 appears to flag defective nascent subunits before they engage in translation. Furthermore, because the arrest of defective subunits could be bypassed by mutations in Tif6 and Nmd3 that facilitate their release from the nascent subunit, we conclude that Tif6 and Nmd3 gate the release of functional subunits into the actively translating pool. How Reh1 collaborates with Tif6 to gate the release of functional subunits remains an open question. However, during the test drive, the P-site ligand Sdo1 and the eEF2-like GTPase Efl1 engage the subunit to evict Tif6 and we have previously shown that mutations in the P-site loop of uL16 impair this step^32^. The N-terminus of Sdo1 inserts into the polypeptide exit tunnel from the PTC and closely approaches the C-terminus of Reh1 which occupies the distal portion of the PET. Consequently, there could be allosteric communication between Sdo1 and Reh1 within the exit tunnel. On the other hand, Reh1 reaches from the exit tunnel to Tif6 on the joining face, raising the possibility that Reh1 could act through stabilizing Tif6 on the joining face. The persistence of either Tif6 or Reh1 could signal to target the defective subunit for degradation.

Previous work to identify quality control factors for defective ribosomes has utilized mutations in ribosomal RNA that lead to rapid decay of defective ribosomes by the nonfunctional RNA Decay pathway^19,20^. Interestingly, the mechanism of degradation of the small subunit is distinct from that of the large subunit. Ribosomes with mutations in the decoding center of the small subunit are recognized during translation elongation and their degradation is initiated through the Integrated Stress Response and involves RNF10-dependent ubiquitination of the subunit^33,51^. In contrast, ribosomes carrying mutations in the PTC of the large subunit appear to enter translation inefficiently^19^. A ubiquitin ligase complex consisting of Mms1, Rtt101 and Crt10 was identified as a major determinant for the accelerated degradation of the 25S NRD pathway^34,52^. The ubiquitinated subunits then appear to be recognized by the ubiquitin binding complex Cdc48-Npl4-Ufd1 for proteasomal degradation^35^. However, it is not yet clear how the defective subunits are initially recognized for targets of ubiquitination. Whether or not Reh1 is a general quality control factor that acts broadly on defective ribosomes, including on these rRNA mutants, remains an open question.

In yeast Reh1 has a paralog, Rei1. However, most eukaryotes have a single gene encoding a Rei1-like protein which is ZNF622 in humans^25^. It is possible that in yeast, a gene duplication event allowed the separation of function of Rei1, which is devoted to ribosome assembly, and Reh1 which has evolved as a quality control factor. During 60S maturation in yeast, Rei1 is replaced by Reh1, which remains on the subunit until Tif6 is released. In human cells, ZNF622 activity is regulated by ubiquitination by the E3 ligase HECTD1 and knocking down HECTD1 leads to increased levels of ZNF622 and stabilization of eIF6 on 60S subunits^53^. A failure to release ZNF622 leads to accumulation of eIF6 which implies that ZNF622 can regulate the release of eIF6 at the licensing step. Consequently, the timing of release of ZNF622 is similar to that of Reh1, raising the interesting possibility that in human cells ZNF622 plays dual roles in 60S maturation and in quality control in human cells.

However, a recent CRISPR screen for ribosome quality control factors in human cells using the equivalent uL16 mutant as we used here, identified the zinc-finger protein ZNF574 as a quality control factor^36^. Knockdown of ZNF574 stabilizes mutant uL16, similar to what we have observed for deletion of *REH1* in yeast. Although how ZNF574 binds to 60S subunits, and how it engages with the nascent subunit to promote turnover is not yet clear. In particular, whether it inserts into the exit tunnel like Reh1 and ZNF622 or binds elsewhere on the subunit to recruit turnover machinery remains to be determined. Additionally, the fact that ZNF622 is codependent on ZNF574 suggests that they are functionally related^36^. Considering the similarities between Reh1 and ZNF622, it is possible that ZNF622 acts as a primary signaling molecule for defective subunits in human cells and that ZNF574 acts downstream of ZNF622 to recruit degradation machinery. Because ZNF622 is required for ribosome assembly^25^, its function as a quality control factor may be masked by its requirement in assembly.

## Materials and Methods

### Yeast strains, plasmids and cell growth

Yeast strains, plasmids and oligonucleotides used in the present study are listed in Supplementary table 1-3. All yeast strains are derivatives of BY4741, unless noted otherwise, and were grown at 30 °C. AJY3918 was made by crossing AJY2846^42^ with an *RPL10* wildtype strain. AJY4618 was made by dissecting the *REH1/reh1Δ* heterozygous diploid (Research Genetics) and converting a resulting spore reh1Δ spore clone to NatMX^54^. AJY4638, AJY4655 and AJY4959 were made by integrating a *TIF6-V192F*:KlURA3 cassette into AJY4618, BY4741 and AJY4638, respectively. The *TIF6- V192F*:KlURA3 cassette was made by overlap PCR of *TIF6* amplified with AJO453xAJO3369 and pFA6a-TAP-KlURA3^55^ amplified with AJO3370xAJO3371. To make AJY4643 the KanMX marker in AJY2104 was converted to NatMX^54^ and then the NatMX:*PGAL-RPL10* locus was amplified and integrated into AJY4638. All plasmids were constructed by Golden gate assembly using the Yeast Tool Kit and BsaI-HF (New England Biolabs) enzyme. *RPL10* alleles were amplified from pAJ1197 or pAJ1777. For pAJ5755 and pAJ5756 a new Yeast Tool Kit Type4a part was generated from pFA6a- ymNeon green-CaURA3 (Addgene plasmid #125703^56^). All plasmid assemblies requiring a PCR-amplified insert were verified via Sanger sequencing. Yeast transformations were all conducted via the poly(ethylene glycol) lithium acetate method. Yeast Toolkit (YTK) plasmids were a gift from J. Dueber (Addgene, kit no. 1000000061).

### Antibodies

The following antibodies were used in this study: Affinity-purified anti-Nmd3^57^ (Dilution 1:5,000), anti-Rpl8 K.-Y. Lo, (Dilution 1:10,000), c-myc 9e10 (626802, Biolegend, Dilution 1:10,000), FLAG M2 (F1804, Sigma, Dilution 1:10,000), and IRDye 680- and 800-labeled secondary antibodies (926-68071 and 926-32210, Dilution 1:15,000–1:20,000).

### Half-life assay

Yeast strains AJY2643 and AJY4638 harboring plasmids pAJ4912 and pAJ4913 were cultured overnight in glucose-based selective medium at 30 °C. Cultures were diluted to OD_600_ = 0.01 in raffinose-containing medium and grown for ∼16 h. The next day, cells were adjusted to OD_600_ = 0.1 in fresh raffinose medium and grown to OD_600_ ∼ 0.3. uL16 expression was induced with 2% galactose for 30 min at 30 °C; an uninduced sample was taken immediately prior to galactose addition. Induction was terminated by adding 2% glucose, and samples were collected at the indicated time points. Cells were harvested by centrifugation and lysed by alkaline extraction. Whole-cell extracts were analyzed by SDS–PAGE and immunoblotting using antibodies against uL16 and G6PD (loading control).

### Sucrose density gradient sedimentation

Yeast cultures were grown continuously on a roller drum overnight to saturation in 10 mL of selective medium, as indicated. The following day, cultures were diluted to an OD_600_of 0.05 in 100 mL of the same medium and grown for 3-4 doublings to reach an OD_600_of 0.4-0.5. Cycloheximide (VWR life sciences) was then added to a final concentration of 100 μg/mL, and cultures were incubated with shaking for an additional 10 min at 30 °C prior to harvesting. Cell pellets were immediately washed and resuspended in 100 μL of lysis buffer containing 20 mM Tris·HCl (pH 7.5), 100 mM NaCl, 30 mM MgCl_2_100 μg/mL CHX, 200 μg/mL heparin, 5 mM β-mercaptoethanol, 1 mM PMSF, and 1 μM each of leupeptin and pepstatin. Cell extracts were prepared by vigorous agitation with glass beads, followed by clarification by centrifugation at 18,000 × g for 15 min at 4 °C. Clarified extracts corresponding to 12 A_260_ units were layered onto 7–47% (w/v) sucrose gradients prepared in TMN buffer (50 mM Tris-acetate, pH 7.0, 50 mM NH_4_Cl, 12 mM MgCl_2_) and centrifuged for 2.5 h at 285,000 × g in a Beckman SW40 rotor. Gradients were fractionated using an ISCO Model 640 fractionator or a Biocomp Piston Gradient Fractionator fitted with a Triax flow cell, collecting 600 μL fractions with continuous absorbance monitoring at 260 nm. To each fraction, 60 μL of 100% trichloroacetic acid (TCA; Sigma T6399) was added, mixed, and incubated at -20 °C overnight. Fractions were centrifuged at 18,000 × g for 15 min at 4 °C, and the resulting protein pellets were washed three times with ice-cold acetone, then resuspended in 1× SDS-PAGE sample buffer and heated at 99 °C for 3 min. Proteins were separated by electrophoresis on 12% SDS-PAGE gels, transferred to nitrocellulose membranes, and analyzed by western blotting using the indicated antibodies (see Antibodies section).

### Immunoprecipitation and Mass Spectrometry (IP–MS)

Cultures of yeast strain AJY2643 and AJY4638 harboring plasmids pAJ5507 or pAJ5509 were grown overnight to saturation in 50 mL of selective medium at 30 °C. The following day, cultures were diluted to an OD_600_ of 0.05 in 500 mL and grown to mid-log phase (OD_600_= 0.4-0.5). Cells were harvested by vacuum filtration using a 0.2 μm aPES membrane bottle-top filter (Fisherbrand), immediately scraped from the filter, and flash-frozen in liquid nitrogen. Frozen cell pellets were stored at -80 °C until further use. For cell lysis, frozen pellets were mixed with frozen IP buffer (20 mM Tris·HCl, pH 7.5, 100 mM KCl, 10 mM MgCl_2_, 100 μg/mL cycloheximide, Pierce Protease Inhibitor Cocktail, EDTA-free [1 tablet/10 mL], 1 mM PMSF, 1 μM each of leupeptin and pepstatin, and 5 mM β-mercaptoethanol) at approximately 1 mL buffer per pellet. Samples were homogenized in a cryogenic mill (Retsch MM 300) for six cycles of 3 min each at 15 Hz, with samples submerged in liquid nitrogen for 3 min between cycles. The resulting cell powder was thawed on ice, additional IP buffer was added, and the lysate was clarified by centrifugation at 18,000 × g for 20 min at 4 °C. The clarified extract was supplemented with 0.1% (v/v) Igepal CA-630.

The lysates were diluted to 150 A260 units/mL. One milliliter of lysate was treated with RNase I (Ambion) at a ratio of 3 U RNase I per 1 A260 unit for 1 h at 4 °C. A 50 μL aliquot of this extract was reserved as the total ribosome profiling sample. The remaining lysate was diluted to 50 A260 units/mL and incubated with 20 μL of pre-washed anti-FLAG magnetic beads (Sigma Millipore) per 1 mL of lysate for 2 h at 4 °C with rotation. Beads were washed three times with IP buffer containing 0.1% Igepal and transferred to a clean tube during the final wash. Bound ribosomes were eluted in 100 μL of IP buffer supplemented with 0.1% Igepal and 150 μg/mL 3×FLAG peptide (Sigma Millipore) for 30 min at 4 °C. Following elution of the uL16 and uL16^mut^ bound ribosomes, samples were subjected to 10–30% (w/v) sucrose density gradient centrifugation prepared in polysome buffer to resolve ribosomal subunits and monosomes. Gradients were centrifuged in a Beckman SW55 rotor at 50,000 rpm for 1.5 h at 4 °C, and fractions corresponding to 60S and 80S ribosomal particles were collected based on absorbance at 260 nm.

Collected fractions were precipitated with TCA, resuspended in SDS sample buffer, and briefly electrophoresed on a 12% Bis-Tris gel (Invitrogen) for approximately 10 min, until the protein mixture formed a single uniform band. The gel region containing the proteins was excised, proteins were digested in-gel with sequencing-grade trypsin digestion and peptides were prepared for LC–MS/MS analysis as previously described^58^. LC-MS/MS data was acquired using an Orbitrap Fusion Tribrid mass spectrometer equipped with an Ultimate 3000 RSLC nano system (Thermo Scientific) with buffer A (0.1% (v/v) formic acid in water) and buffer B (0.1% (v/v) formic acid in acetonitrile). Peptides were concentrated onto an PepMap Neo C18 Trap column (5 um 300 μm x 5 mm, Thermo Scientific) then separated on a 75 μm × 50 com C18 column (PepMap RSLC, Thermo Scientific). Liquid chromatography was performed using a gradient of 3–35% buffer B over 57 min.

The resulting spectra were processed with Proteome Discoverer version 2.5 (Thermo Scientific) using the Sequest HT search engine, with 10 ppm mass tolerance for the MS at 0.6 Da for the MS/MS. Identifications were validated with Scaffold 5 (Proteome Software). Raw data were processed with MaxQuant^59^ using default parameters to obtain label-free quantification (LFQ) intensities. Reverse hits, contaminants, and site-only identifications were removed. Protein intensities were log_2_-transformed, and missing values were imputed using the K-nearest neighbors algorithm (impute R package, v1.82.0) to approximate low-abundance protein detection. Differential abundance analysis between control and treatment samples was performed in R (v4.3.1) using the limma package (v3.64.3)^60^. Proteins with log₂ fold change > 0.9 and FDR < 0.05 (Benjamini–Hochberg correction) were considered significant. Volcano plots were generated using ggplot2 and ggrepel to visualize differential protein expression.

### Ribosome Profiling experiments and computational analysis

Immunoprecipitation was performed as described for Immunoprecipitation and Mass Spectrometry using extracts from *S. cerevisiae* strain AJY4638 harboring plasmid pAJ5507 with a modified IP buffer (20 mM Tris·HCl, pH 7.5, 100 mM KCl, 10 mM MgCl2, 100 μg/mL cycloheximide, 100 μg/mL Tigecyclin (Sigma), Pierce Protease Inhibitor Cocktail, EDTA-free [1 tablet/10 mL], 1 mM PMSF, 1 μM each of leupeptin and pepstatin, and 5 mM β-mercaptoethanol). Eluted ribosomes were digested with RNase I (1 U/μg RNA; Ambion) for 45 min at room temperature. Reactions were stopped by addition of QIAzol (Qiagen), and RNA was extracted using chloroform extraction followed by ethanol precipitation. Total ribosome profiling samples were processed in parallel by treating 50 μL of lysate with RNase I (1 U/μg RNA) in IP buffer supplemented with 0.1% Igepal for 45 min at room temperature. Digested lysates were layered onto 50 μL of 1 M sucrose cushions prepared in IP buffer, and ribosomes were pelleted by centrifugation at 165,000 × g for 30 min at 4 °C in a TLA 100 rotor (Beckman Coulter).

RNA was extracted from ribosome pellets using 700 μL of QIAzol, followed by chloroform extraction and ethanol precipitation. To isolate the ribosome-protected fragments (RPFs), the digested RNA was run on a 15% TBE–urea polyacrylamide gel (Invitrogen). A 19– 35 nt fragments were excised and extracted by crushing the excised gel slice in 400 μL RNA extraction buffer (300 mM sodium acetate, pH 5.5, 5 mM MgCl2 followed by incubation at -20 C for 1h and overnight incubation at RT on a hula mixer. The gel extract was passed through a Spin-X filter (Corning 8160), and RNA was recovered from the flow-through by ethanol precipitation (2.5X volume) with 1.5 μL GlycoBlue (Invitrogen AM9516).

Ribosome profiling libraries were prepared as previously described^61^ using the D-Plex Small RNA-Seq Kit (Diagenode). cDNA was subjected to rRNA depletion, and the cleaned-up cDNA was amplified by PCR for 12 cycles. Libraries were purified with AMPure XP (Beckman Coulter) at a ratio of 1:1.8 (sample to bead ratio) and eluted in 50 μL of water. The library quality was assessed using a Bioanalyzer and the Agilent High Sensitivity DNA Kit (Agilent). Final libraries were pooled in equimolar ratios and sequenced on the NovaSeqX Plus; PE 150 (Illumina).

Ribosome profiling data was preprocessed using RiboFlow^62^. Briefly, the first 12 nucleotides were extracted and used as unique molecular identifiers. The next four nucleotides matching the sequence ‘NGGG’ were discarded as these were added during the template-switching reverse transcription step. Sequencing fragments containing the 3’ adapter sequence AAAAAAAAAACAAAAAAAAAA were retained for further analyses. rRNA, tRNA and mRNA sequences were obtained from SGD on March 14, 2022. Sequences for the CDS, 5’ UTR (untranslated region) and 3’ UTR were merged to form the complete mRNA transcript giving us a total of 5024 transcripts. For transcripts annotated with more than one 3’ UTR sequence, we randomly retained one of the sequences. The resulting reference sequences are available at https://github.com/ribosomeprofiling/yeast_reference. PCR duplicates were eliminated from the ribosome profiling data using UMI-tools^63^. Initial data QC was visualized using RiboGraph^64^ and statistical analysis and meta-gene plots were generated using RiboR with ribosome footprints of length between 26-29 and separately for 30 nt^62^. The start site ratio was calculated as the sum of reads at the start codon divided by CDS reads for each transcript, and log₂ fold changes between IP and total ribosome samples were visualized as histogram.

### Cryo-EM sample preparation, data collection, and image processing

Immunoprecipitation was performed as described above using extracts from yeast strain AJY2643 and AJY4638 harboring plasmid pAJ5507. Eluates were either frozen or else directly applied to cryo-EM grids. Quantifoil R1.2/1.3, 200-mesh copper grids coated with an ultrathin layer of amorphous carbon were plasma cleaned for 30 s at 25 mA using a Solarus 950 plasma cleaner (Gatan). A 2.5 μL aliquot of sample was applied to a freshly cleaned grid, blotted for 6 s with a blot force of 0 in a Vitrobot Mark IV (Thermo Fisher Scientific) at 4 °C and 100% humidity, and plunge-frozen in liquid ethane. Grids were stored in liquid nitrogen before screening in SerialEM^65^ on either a Glacios cryo-TEM or an FEI Titan Krios cryo-TEM. For samples derived from wild-type cells, data were collected on a Glacios cryo-TEM equipped with a Falcon 4 direct electron detector at a pixel size of 0.933 Å. For samples from the bypass mutant, data were collected on an FEI Titan Krios cryo-TEM equipped with a Gatan K3 direct electron detector at a pixel size of 0.8332 Å. The defocus range for all datasets was −1.0 to −2.0 μm. Motion correction, contrast transfer function (CTF) estimation, and blob-based particle picking were performed in cryoSPARC Live v4.0, and all subsequent processing steps were carried out in cryoSPARC v4.4^66^. For the wild-type dataset, 158,256 particles were picked from 2,199 accepted micrographs and subjected to 2D classification. A subset of 129,471 particles from selected 2D classes was used for ab initio reconstruction into three classes, followed by one round of heterogeneous refinement. The best-resolved class was re-extracted at a box size of 600 pixels, then processed by global and local CTF refinement and non-uniform refinement, yielding a 2.73 Å reconstruction from 76,050 particles (Supplementary Fig. 1). To assess Yvh1 occupancy, these particles were subjected to focused 3D classification using a molmap mask around Yvh1 generated in ChimeraX (v1.9)^67^. The resulting classes were further sorted into four classes based on Yvh1 occupancy (Supplementary Fig. 1). The Yvh1-bound class yielded a 2.96 Å reconstruction from 27,593 particles (Supplementary Fig. 3a).

For the bypass mutant dataset, 898,318 particles were initially picked. After 2D classification, 597,679 particles were retained for ab initio reconstruction into three classes, followed by heterogeneous refinement, non-uniform refinement and re-extraction at a box size of 600 pixels. This yielded a 60S subunit reconstruction from 204,666 particles and an 80S subunit reconstruction from 197,127 particles. Focused 3D classification for the 60S subunit was performed using a molmap mask around NMD3 generated in ChimeraX and yielded two maps: a 2.78 Å reconstruction lacking LSG1, NMD3, but containing TIF6, and a 2.83 Å reconstruction containing all three proteins (Supplementary Fig. 2, Supplementary Fig. 3b-c). To examine tRNA heterogeneity in the 80S subunit, particles were subjected to 3D classification, producing four distinct tRNA-binding states: (i) P/E and A/P tRNA, (ii) A-site and P-site tRNA, (iii) P-site tRNA only, and (iv) tRNA-free 80S subunit (Supplementary Fig. 2, Supplementary Fig. 4a-c). To assess the density within the exit tunnel, focused classification was performed using a molmap-derived mask encompassing the tunnel region, which identified particles containing Reh1 density in the exit tunnel with varying tRNA occupancies (Supplementary Fig. 2, Supplementary Fig. 4d-e).

### Model building and refinement

Atomic models for the 60S and 80S subunits were initiated from PDB entries 6N8O^24^ and 8UT0^40^, respectively. Manual model adjustment was performed in Coot (v0.9.8.95)^68^, followed by automated refinement using the real_space_refine in PHENIX (v1.21.2- 5419)^69^. All structural figures were prepared in ChimeraX (v1.9).

### Correlation and Gene Ontology Analysis

Proportionality metrics rho (ρ) from Rich et al.^47^ were used to rank genes by co-expression with Reh1 or Rei1. A pre-ranked GO enrichment analysis was performed using fgsea (v 1.34.2)^70^ with annotations from the Saccharomyces Genome Database, filtering GO terms with 5-500 genes. Top 10 significantly enriched GO terms (adjusted p < 0.05) were visualized as violin plots showing the distribution of proportionality values for highly and lowly co-expressed gene sets. To compare Reh1 with other ribosome biogenesis, quality control, and proteasomal factors, correlation scores were summarized by median values across selected GO categories and visualized as a clustered heatmap in R.

## Acknowledgements

This work was supported by NIH grants GM127127 (to A.W.J.) GM138348 (to D.W.T). and GM150667 (to CC), and Welch Foundation Research Grants F-1938 (to D.W.T) and F-2027-20230405 (to C.C.). We thank D. Gottschling for pAGL, J. Tesmer for pMALC2H10T, B. Bass for pMP004, J. Dueber for yeast toolkit vectors and K.-Y. Lo for rabbit anti-Rpl8 and anti-Tif6 antibody and J. Black for initial analysis of mutant uL16 in the bypass strain. Protein identification was provided by the UT Austin Center for Biomedical Research Support Biological Mass Spectrometry Facility (RRID:SCR_021728).

## Author Contributions

R.C. conceived the experiments, performed genetic and biochemical analysis, analyzed the ribosome profiling data, prepared samples for cryo-EM and wrote the manuscript.

K.G. performed cryo-electron microscopy, structure determination and modeling and wrote the manuscript. S.R. performed ribosome profiling experiments. C.W. performed western blotting for protein stability determination. C.C. supervised the bioinformatic analyses and obtained funding for the work. D.W.T. supervised the structural studies and obtained funding for the work. A.W.J. conceived the experiments, performed genetic and biochemical analysis, analyzed the data and obtained funding for the work. All authors contributed to editing the manuscript.

## Data availability

Cryo-EM map of uL16^mut^ pre-60S ribosomes produced in WT strain have been deposited in the Electron Microscopy Data Bank (EMDB) under accession code EMD-72796. Maps of uL16^mut^ pre-60S ribosomes produced in bypass strain without or with Lsg1, Nmd3 have been deposited as EMD-72797 and EMD-72799, respectively. The corresponding atomic model for EMD-72799 has been deposited to the Protein Data Bank (PDB) under accession code PDB-9YDB. Cryo-EM maps of 80S ribosome containing A/P P/E tRNA, A- P-site tRNA and P-site tRNA have been deposited as EMD-72800, EMD-72801 and EMD-72802, respectively, with the corresponding models deposited as PDB-9YDC, 9YDD, and 9YDE. Cryo-EM maps of 80S ribosome containing Reh1 in exit tunnel, either without tRNA or with a P-site tRNA, have been deposited as EMD-72803 and EMD- 72804, respectively. Sequencing files for selective ribosome profiling experiments will be available at GEO (accession number GSEXXX).

