## Supplemental Materials for "A late cytoplasmic surveillance pathway ensures ribosome integrity"

**Supplementary Table 1:**

| Strain | Genotype | Source |
|---|---|---|
| AJY2104 | *MATalpha KanMX:P$_{GAL}$ RPL10 ade2 ade3 ura3 leu2* | [1] |
| AJY3918 | *MATalpha NMD3-Y379D his3Δ1 leu2Δ0 ura3Δ0 met15Δ* | This work |
| AJY4618 | *MATa reh1Δ::NatMX his3Δ1 leu2Δ0 ura3Δ0 met15Δ* | This work |
| AJY4638 | *MATalpha NMD3-Y379D TIF6-V192F:KlURA3 his3Δ1 leu2Δ0 met15Δ* | This work |
| AJY4643 | *MATalpha NMD3-Y379D TIF6-V192F:KlURA3 NatMX:P$_{GAL}$ RPL10 his3Δ1 leu2Δ0 met15Δ* | This work |
| AJY4655 | *MATalpha TIF6-V192F:KlURA3 his3Δ1 leu2Δ0 met15Δ* | This work |
| AJY4959 | *MATalpha reh1Δ::KanMx NMD3-Y379D TIF6-V192F:KlURA3 his3Δ1 leu2Δ0 met15Δ* | This work |
| BY4741 | *MATa his3Δ1 leu2Δ0 ura3Δ0 met15Δ* | |

**Supplementary Table 2:**

| Plasmid | Features | Source |
|---|---|---|
| pAJ1197 | *P$_{RPL10}$-RPL10-13myc LEU2* CEN Ampr | [1] |
| pAJ1777 | *P$_{RPL10}$-rpl10Δ102-112-13xmyc LEU2* CEN Ampr | [1] |
| pAJ4796 | *P$_{RPL18B}$-RPL10-13myc LEU2* CEN | This work |
| pAJ4797 | *P$_{RPL18B}$-rpl10Δ102-112-13xmyc LEU2* CEN Ampr | This work |
| pAJ4912 | *P$_{GAL}$-RPL10-3xFLAG LEU2* CEN Ampr | This work |
| pAJ4913 | *P$_{GAL}$-rpl10Δ102-112-3xFLAG LEU2* CEN Ampr | This work |
| pAJ5507 | *P$_{RPL18B}$-rpl10Δ102-112-3xFLAG LEU2* CEN Ampr | This work |
| pAJ5509 | *P$_{RPL18B}$-RPL10-3xFLAG LEU2* CEN Ampr | This work |
| pAJ5755 | *P$_{RPL18B}$-RPL10-ymNeonGreen LEU2* CEN Amp | This work |
| pAJ5756 | *P$_{RPL18B}$-rpl10Δ102-112-ymNeonGreen LEU2* CEN Ampr | This work |
| pRS315 | *LEU2* CEN Ampr | This work |

**Supplementary Table 3:**

| Oligos | Sequence | Source |
|---|---|---|
| AJO264 | 5'- CGCGGAtccgaaactagttagcac | [1] |
| AJO453 | 5'- cggaagcttGCATTCTGGACGAAATCC | This work |
| AJO754 | 5'- cgcGGATCCgaattcatgaacgggaaa | This work |
| AJO3369 | 5'- CTATGAGTAGGTTTCAATCAAAGTATCACG | This work |
| AJO3370 | 5'- CGTGATACTTTGATTGAAACCTACTCATAGggcgcgc ctctctgc | This work |
| AJO3371 | 5'-TGTAAACAGACTTGAGGAAGGAGGGGAATCCCCTC AGGAGTACCTGACATgtttgagagggcttatcgc | This work |
| AJO4016 | 5'- ggtggtctcatATGGCTAGAAGACCAGCTAG | This work |
| AJO4041 | 5'- ggcGGTCTCAGGATGCTTGAGCAGCAAAGTATTCT | This work |
| AJO4507 | 5'- tagcCGTCTCATCGGTCTCAATCCTCTGTCTCTAA | This work |
| AJO4508 | 5'- tagcCGTCTCAGGTCTCAGCCATTACTTGTACAAT | This work |

### Supplementary Table 4. Cryo-EM data collection and model validation statistics

| | 60S (WT) | 60S (bypass) Lsg1, Nmd3 absent, Tif6 presents. | 60S (bypass) Lsg1, Nmd3, Tif6 present. | 80S (bypass) A/P, P/E tRNA | 80S (bypass) A- and P-site tRNA | 80S (bypass) P-site tRNA | 80S (bypass) tRNA absent, Reh1 in exit tunnel | 80S (bypass) P-site tRNA, Reh1 in exit tunnel |
|---|---|---|---|---|---|---|---|---|
| **Data collection and processing** | | | | | | | | |
| Magnification | 150K | 105K | 105K | 105K | 105K | 105K | 105K | 105K |
| Voltage (kV) | 200 | 300 | 300 | 300 | 300 | 300 | 300 | 300 |
| Electron exposure (e–/Å$^2$) | 50 | 80 | 80 | 80 | 80 | 80 | 80 | 80 |
| Defocus range (μm) | -1.0 to -2.0 | -1.0 to -2.0 | -1.0 to -2.0 | -1.0 to -2.0 | -1.0 to -2.0 | -1.0 to -2.0 | -1.0 to -2.0 | -1.0 to -2.0 |
| Pixel size (Å) | 0.94 | 0.83 | 0.83 | 0.83 | 0.83 | 0.83 | 0.83 | 0.83 |
| Initial particle (no.) | 76, 050 | 203, 066 | 203, 066 | 187, 670 | 187, 670 | 187, 670 | 187, 670 | 187, 670 |
| Final particle (no.) | 27, 593 | 103, 394 | 99, 672 | 15, 817 | 79, 307 | 35, 555 | 34, 116 | 6, 914 |
| Map resolution (Å) | 2.96 | 2.78 | 2.83 | 3.02 | 2.66 | 2.82 | 3.01 | 3.45 |
| FSC threshold | 0.143 | 0.143 | 0.143 | 0.143 | 0.143 | 0.143 | 0.143 | 0.143 |
| **Refinement** | | | | | | | | |
| Initial model used | - | - | 6N8O | 6N8O/8UTI | 6N8O/8UTI | 6N8O/8UTI | - | - |
| Model resolution (Å) | - | - | 3.0 | 3.2 | 2.9 | 3.0 | - | - |
| FSC threshold | - | - | 0.5 | 0.5 | 0.5 | 0.5 | - | - |
| Map sharpening B factor (Å$^2$) | -26.0 | -47.1 | -43.7 | -18.6 | -43.1 | -33.5 | -35.7 | -4.9 |
| Model composition | | | | | | | | |
| Non-hydrogen atoms | - | - | 126, 907 | 194, 693 | 194, 353 | 192, 546 | - | - |
| Protein residues | - | - | 7,082 | 10, 896 | 10, 916 | 10, 891 | - | - |
| Nucleotides | - | - | 3,302 | 5, 079 | 5, 056 | 4979 | - | - |
| Ligands | - | - | 3HE: 1 | - | 3HE: 1 | 3HE: 1 | - | - |
| Mean B factors (Å$^2$) | - | - | | | | | - | - |
| Protein | - | - | 83.01 | 96.95 | 81.63 | 95.32 | - | - |
| Nucleotides | - | - | 76.37 | 98.45 | 75.65 | 93.62 | - | - |
| Ligands | - | - | 55.13 | - | 46.84 | 58.64 | - | - |
| R.m.s deviations | - | - | | | | | - | - |
| Bond lengths (Å) | - | - | 0.005 | 0.004 | 0.006 | 0.005 | - | - |
| Bond angles (°) | - | - | 0.939 | 0.929 | 0.948 | 0.937 | - | - |
| **Validation** | - | - | | | | | - | - |
| MolProbity score | - | - | 1.65 | 1.61 | 1.65 | 1.61 | - | - |
| Clashscore | - | - | 4.24 | 4.35 | 4.26 | 4.25 | - | - |
| Poor rotamers (%) | - | - | 2.18 | 1.58 | 2.12 | 1.87 | - | - |
| Ramachandran plot | - | - | | | | | - | - |
| Favored (%) | - | - | 96.88 | 96.33 | 96.82 | 96.81 | - | - |
| Allowed (%) | - | - | 3.10 | 3.57 | 3.16 | 3.10 | - | - |
| Disallowed (%) | - | - | 0.01 | 0.10 | 0.02 | 0.09 | - | - |
| **PDB** | **-** | **-** | **9YDB** | **9YDC** | **9YDD** | **9YDE** | **-** | **-** |
| **EMDB** | **EMD-72796** | **EMD-72797** | **EMD-72799** | **EMD-72800** | **EMD-72801** | **EMD-72802** | **EMD-72803** | **EMD-72804** |

**Supplementary Figures:**

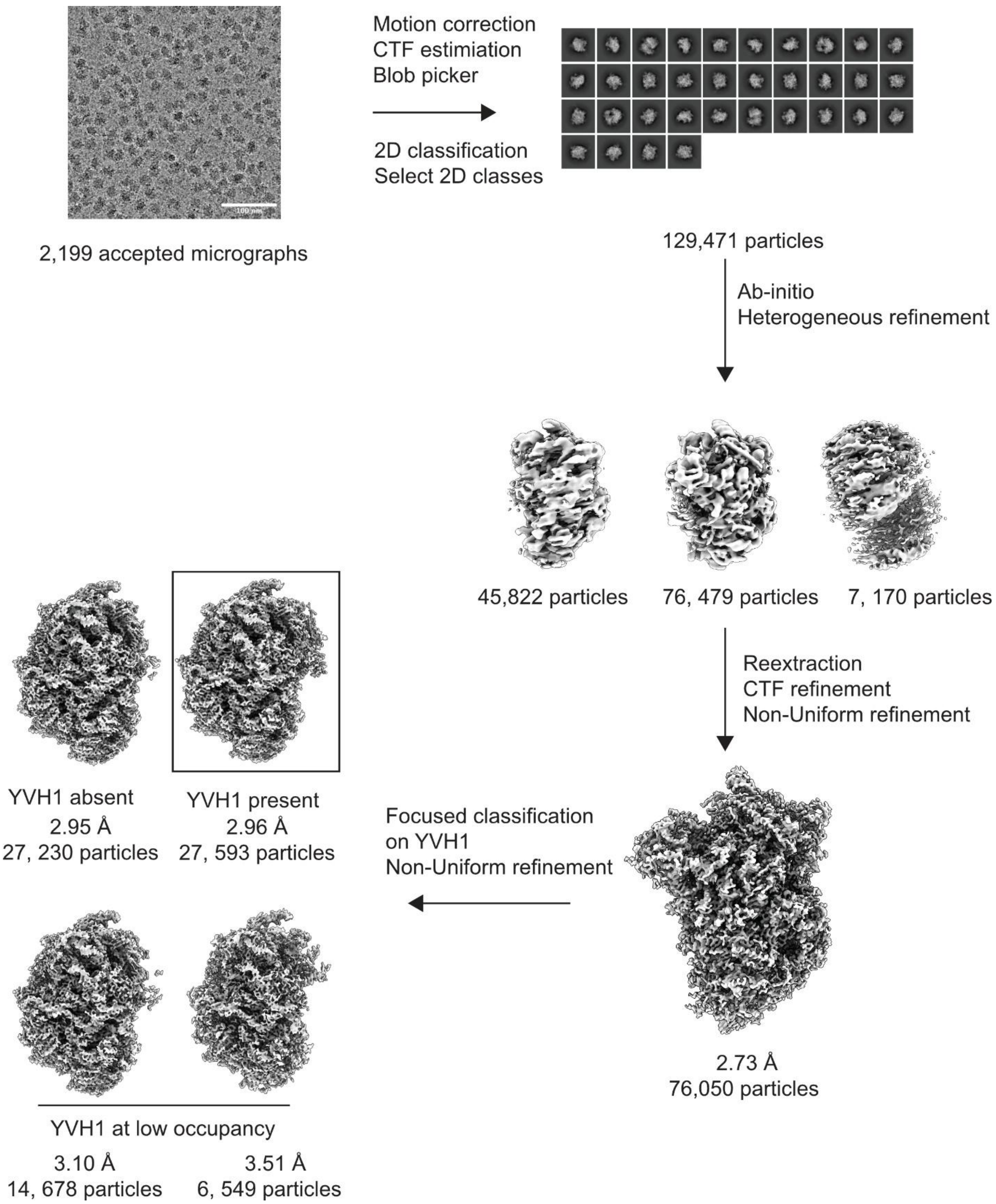


**Supplementary Figure 1**: Cryo-EM data processing workflow for uL16$^{mut}$ sample produced in WT strain. Maps in box were presented and deposited.

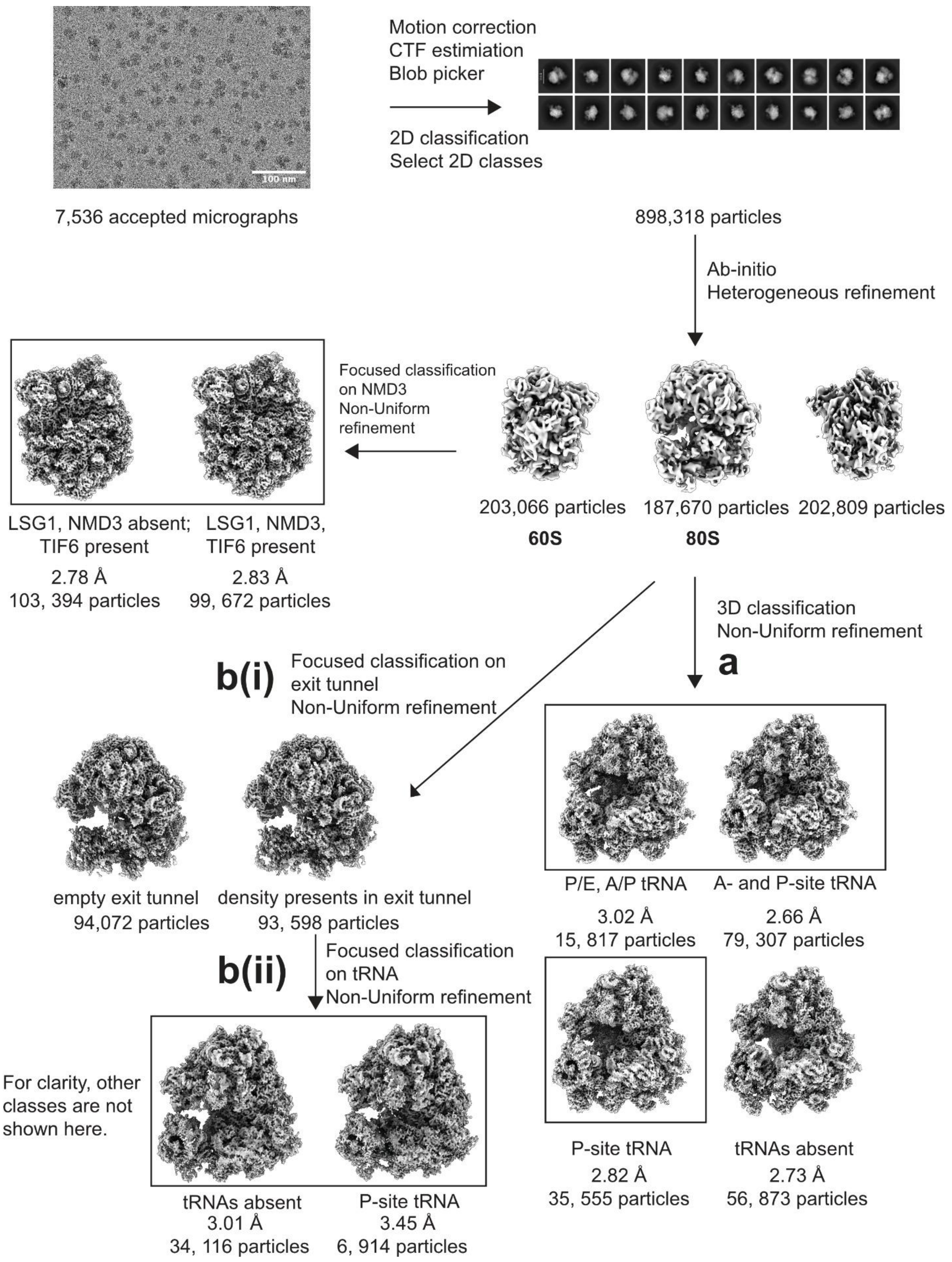

Motion correction
CTF estimiation
Blob picker
2D classification
Select 2D classes
100 nm
7,536 accepted micrographs
898,318 particles
Ab-initio
Heterogeneous refinement
Focused classification
on NMD3
Non-Uniform
refinement
203,066 particles
60S
187,670 particles
80S
202,809 particles
LSG1, NMD3 absent;
TIF6 present
2.78 Å
103, 394 particles
LSG1, NMD3,
TIF6 present
2.83 Å
99, 672 particles
3D classification
Non-Uniform refinement
a
b(i)
Focused classification on
exit tunnel
Non-Uniform refinement
empty exit tunnel
94,072 particles
density presents in exit tunnel
93, 598 particles
P/E, A/P tRNA
3.02 Å
15, 817 particles
A- and P-site tRNA
2.66 Å
79, 307 particles
b(ii)
Focused classification
on tRNA
Non-Uniform refinement
For clarity, other
classes are not
shown here.
tRNAs absent
3.01 Å
34, 116 particles
P-site tRNA
3.45 Å
6, 914 particles
P-site tRNA
2.82 Å
35, 555 particles
tRNAs absent
2.73 Å
56, 873 particles

**Supplementary Figure 2:** Cryo-EM data processing workflow for uL16$^{mut}$ sample produced in bypass mutant strain. Maps in box were presented and deposited. (a), (b) are classification strategies. **(a)** 3D classification based on tRNAs. **(b)(i)** Focused classification on exit tunnel. **(b)(ii)** Particles showing density in exit tunnel are further classified based on tRNAs.

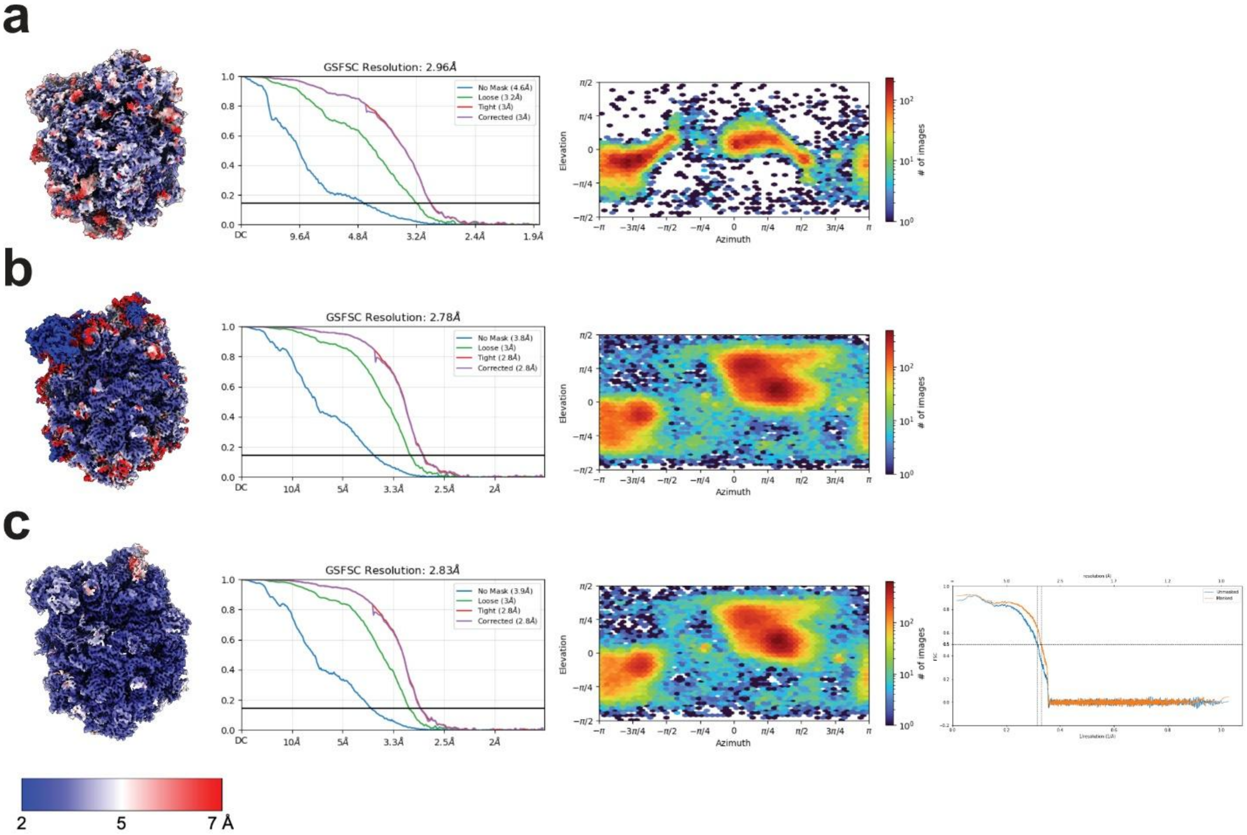


**Supplementary Figure 3:** Cryo-EM data analysis. **(a)** 60S from WT strain with Yvh1 present; **(b)** 60S from bypass strain with Lsg1, Nmd3 and Tif6 absent and Reh1 present; **(c)** 60S from bypass strain with Lsg1, Nmd3, Tif6 and Reh1 present. For each structure, left to right panels show: local resolution map; gold-standard Fourier Shell Correlation (FSC) curves with resolution reported at FSC= 0.143; Euler angle distribution plots of particle orientations; map-to-model FSC curves.

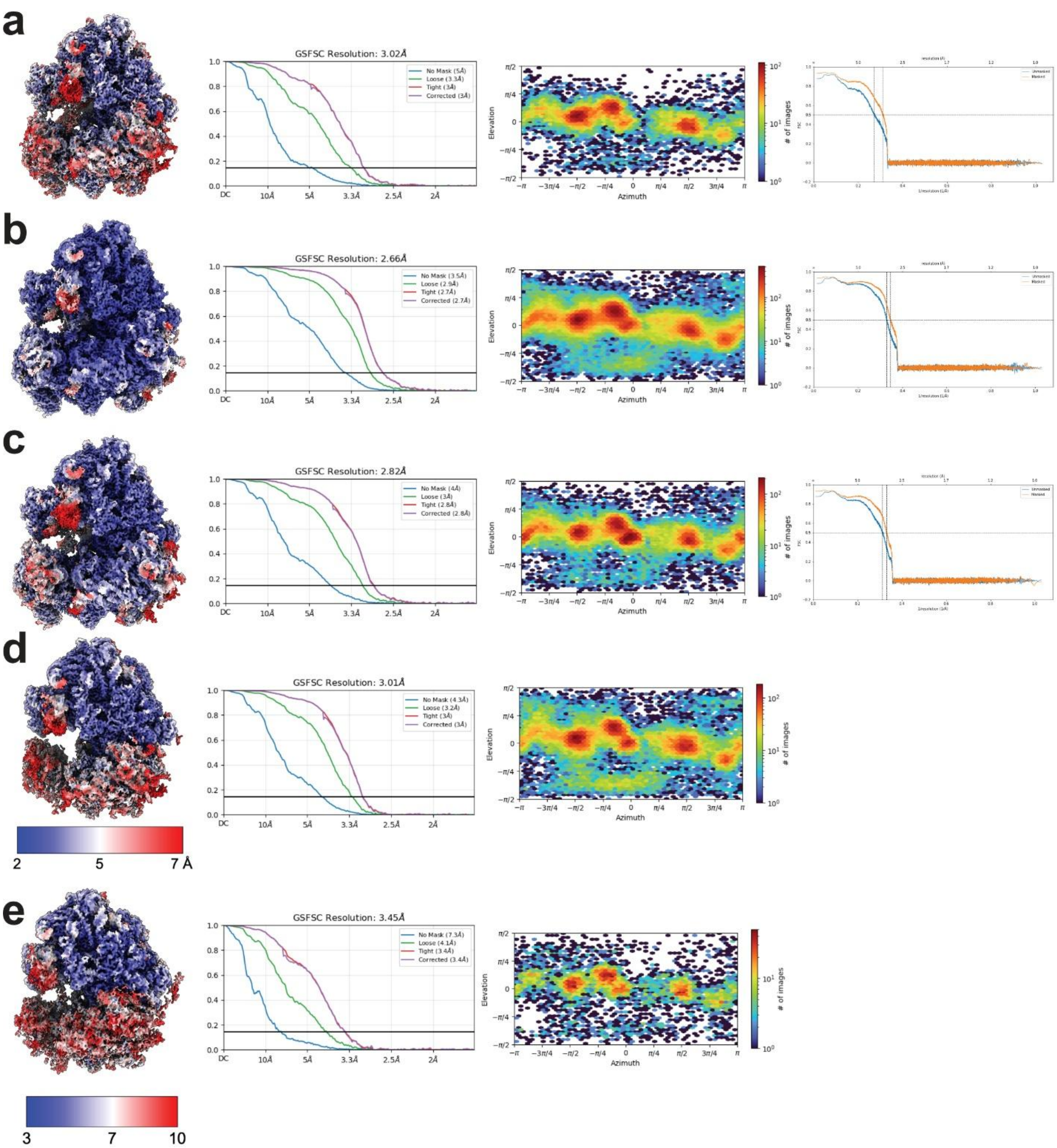


**Supplementary Figure 4:** Cryo-EM data analysis. **(a)** 80S from bypass strain, P/E and A/P tRNA; **(b)** 80S from bypass strain, P- and A-site tRNA; **(c)** 80S from bypass strain, P-site tRNA; **(d)** 80S from bypass strain, tRNAs absent; Reh1 in exit tunnel. **(e)** 80S from bypass strain, P-site tRNA; Reh1 in exit tunnel. For each structure, left to right panels show: local resolution map; gold-standard Fourier Shell Correlation (FSC) curves with resolution reported at FSC= 0.143; Euler angle distribution plots of particle orientations; map-to-model FSC curves.

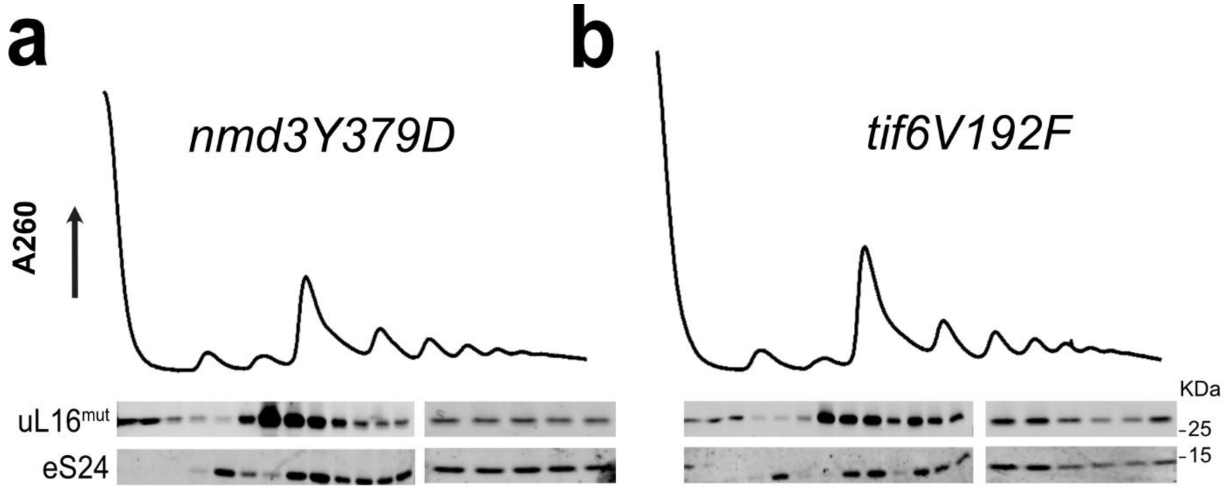


**Supplementary Figure 5:** Sucrose density gradient sedimentation of plasmid-borne uL16$^{mut}$ in *nmd3-Y379D* **(a)** and *tif6-V192F* **(b)** single mutants. UV trace monitoring A260 is shown. Fractions were analyzed by western blotting for the presence uL16$^{mut}$ -FLAG, and eS24.

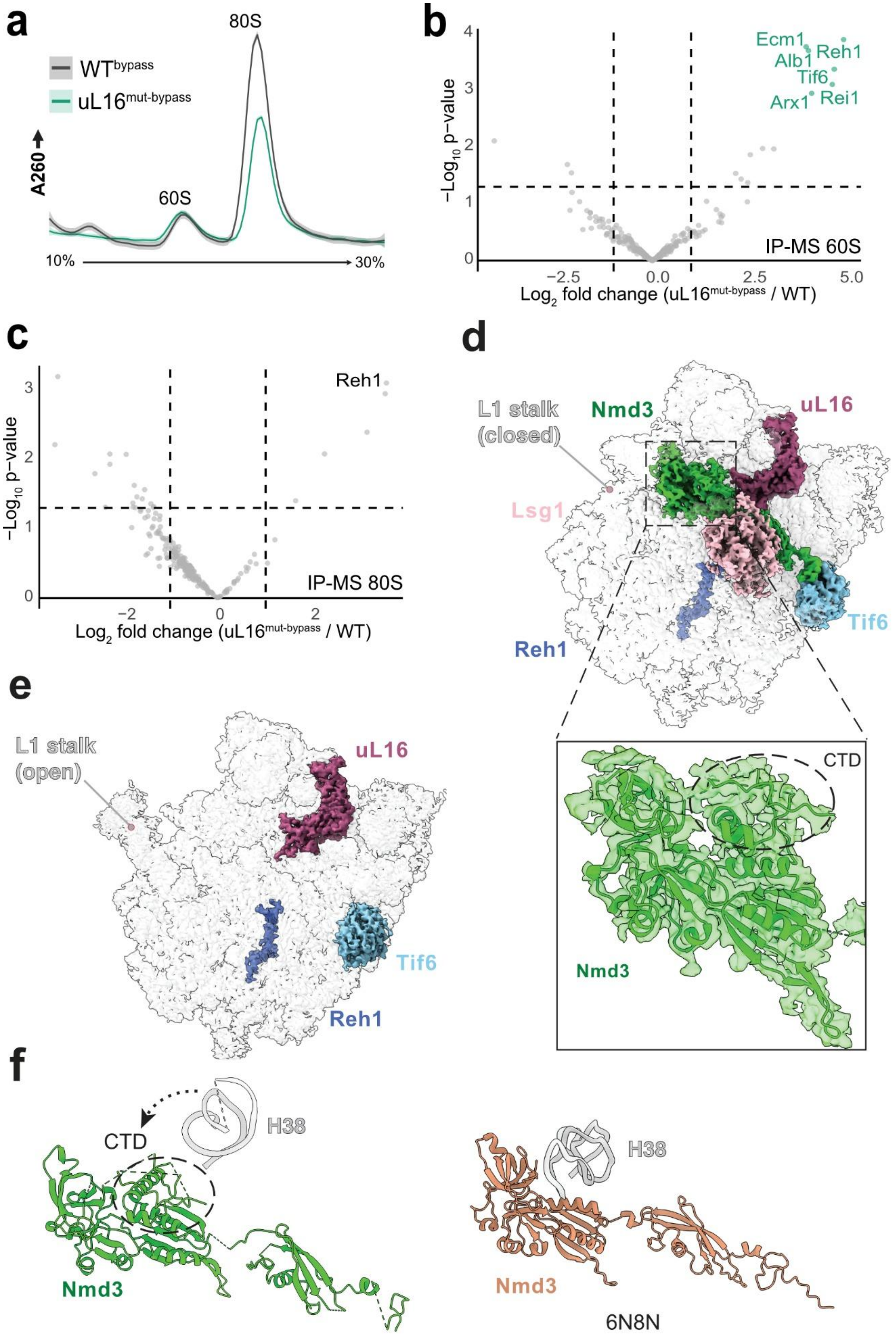
a
80S
WT^bypass
uL16^mut-bypass
A260
60S
10%
30%
b
Ecm1
Alb1
Reh1
Tif6
Arx1
Rei1
−Log10 p−value
IP-MS 60S
Log2 fold change (uL16^mut-bypass / WT)
c
Reh1
−Log10 p−value
IP-MS 80S
Log2 fold change (uL16^mut-bypass / WT)
d
L1 stalk (closed)
Nmd3
uL16
Lsg1
Tif6
Reh1
CTD
Nmd3
e
L1 stalk (open)
uL16
Tif6
Reh1
f
CTD
H38
Nmd3
H38
Nmd3
6N8N

**Supplementary Figure 6: Cryo-EM analysis of uL16$^{mut}$ 60S subunits from the bypass strain reveals partial suppression of the biogenesis arrest. (a)** The uL16$^{mut}$ associates with 60S and 80S particles in the bypass strain. FLAG-tagged wild-type and uL16$^{mut}$ were expressed in *nmd3-Y379D tif6-V192f* cells and affinity-purified complexes were analyzed on sucrose density gradients. UV absorbance at A260 is shown for purified ribosomal complexes. **(b)** Volcano plot comparing LC-MS/MS results from 60S fractions of affinity-purified FLAG-tagged wild-type and uL16$^{mut}$ -containing ribosomes from (a). Fold changes and p-values (Student's t-test) are shown; proteins of interest are highlighted. **(c)** Volcano plot comparing LC-MS/MS results from 80S fractions of affinity-purified FLAG-tagged wild-type and uL16$^{mut}$-containing ribosomes from (a). Fold changes and p-values (Student's t-test) are shown; proteins of interest are highlighted. **(d)** Cryo-EM map of a pre-60S particle purified from uL16$^{mut}$ ribosomes in the bypass strain, showing associated biogenesis factors Nmd3, Tif6, Lsg1, and Reh1. **(e)** Cryo-EM map of a distinct pre-60S particle from the same mutant strain showing only Tif6 and Reh1, indicative of partial release of biogenesis factors and bypass of arrest. **(f)** The CTD of Nmd3 is absent from structures where helix 38 (H38) of the 25S rRNA is engaged with the eIF5A domain of Nmd3 (right, PDB:6N8N). Upon retraction of H38, the Nmd3 CTD becomes visible (left).

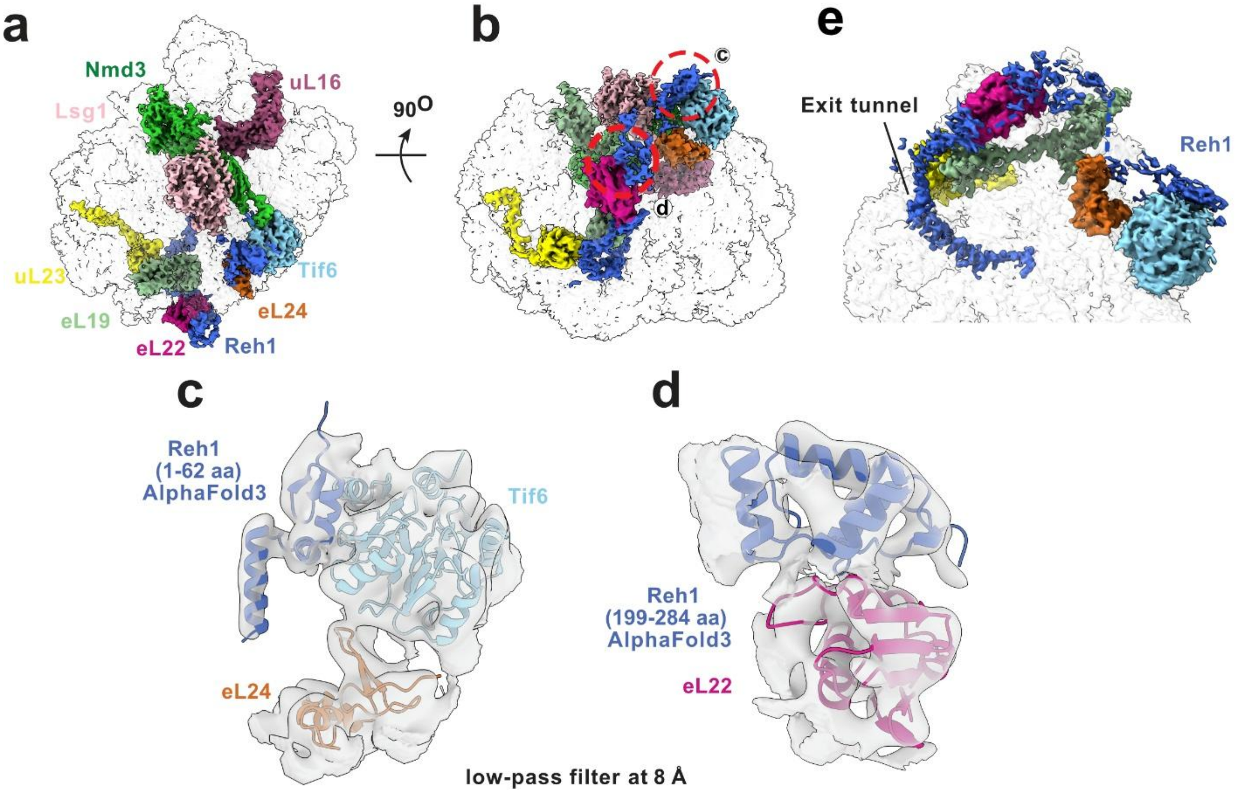


**Supplementary Figure 7: Interaction sites of Reh1 on the 60S ribosomal subunit. (a)** Cryo-EM map of a pre-60S particle from the bypass strain showing associated biogenesis factors and ribosomal proteins. Reh1 density is shown in blue. The first zinc cluster of Reh1 (residues 6–30) along with a α-helix pass in close proximity to Tif6 and eL24. **(b)** A disordered region of Reh1 (residues 72–150) is observed traversing along an rRNA element. The second (residues 186–209) and third (residues 237–261) zinc clusters of Reh1 contact eL22. **(c)** Low-pass filtered Cryo-EM map of the pre-60S particle overlaid with an AlphaFold-predicted model of the Reh1 N-terminus (green), showing a good fit of the first zinc cluster interacting with Tif6. **(d)** AlphaFold model of the second and third zinc clusters of Reh1 (residues 199–284) fits well into the low-pass filtered Cryo-EM map and is observed interacting with eL22. **(e)** On the solvent-exposed surface of the pre-60S, Reh1 density is observed in proximity to eL19 and uL23 and is seen extending into the ribosomal exit tunnel.

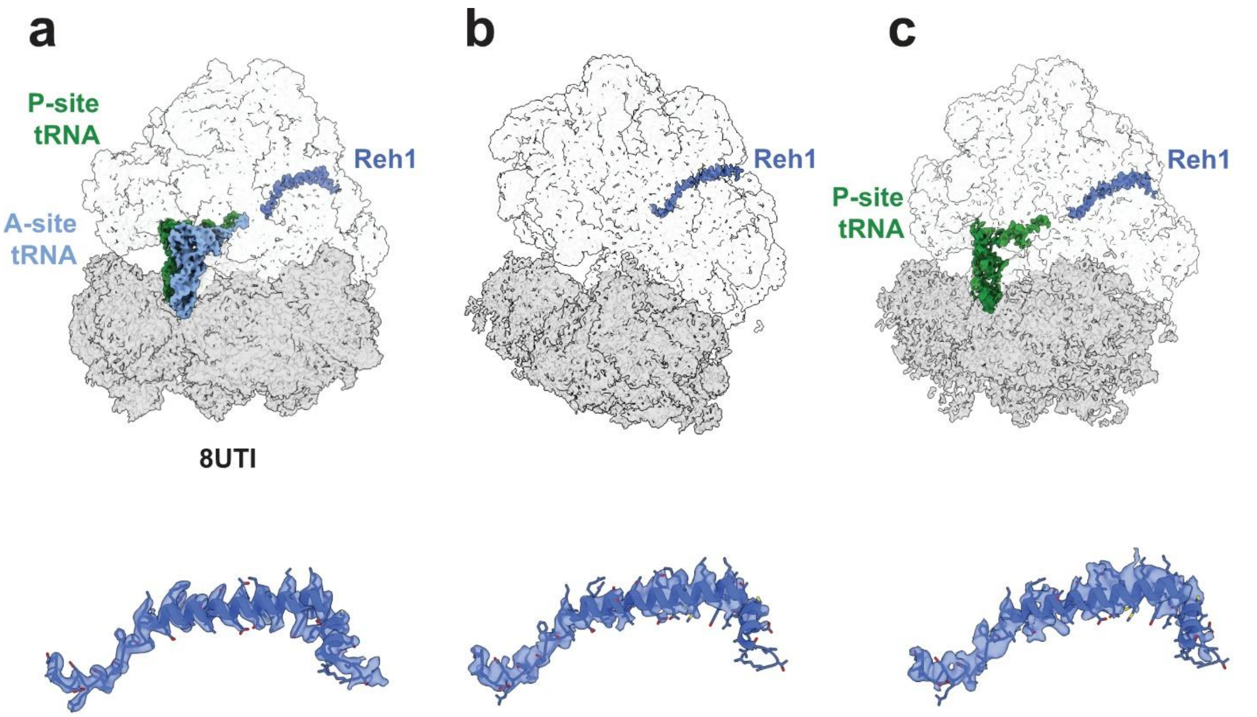


**Supplementary Figure 8. Reh1 is retained in the exit tunnel of 80S ribosomes from the uL16$^{mut}$ in the bypass strain. (a**) Fitted model of Reh1 within the cryo-EM density of 80S ribosomes carrying classical A- and P-site tRNAs (PDB: 8UTI). **(b)** Fitted Reh1 model within the cryo-EM density of empty 80S ribosomes derived from the bypass strain. **(c)** Fitted Reh1 model within the cryo-EM density of 80S ribosomes carrying a P-site tRNA derived from the bypass strain. Reh1 adopts distinct conformations in the exit tunnel in these structures.

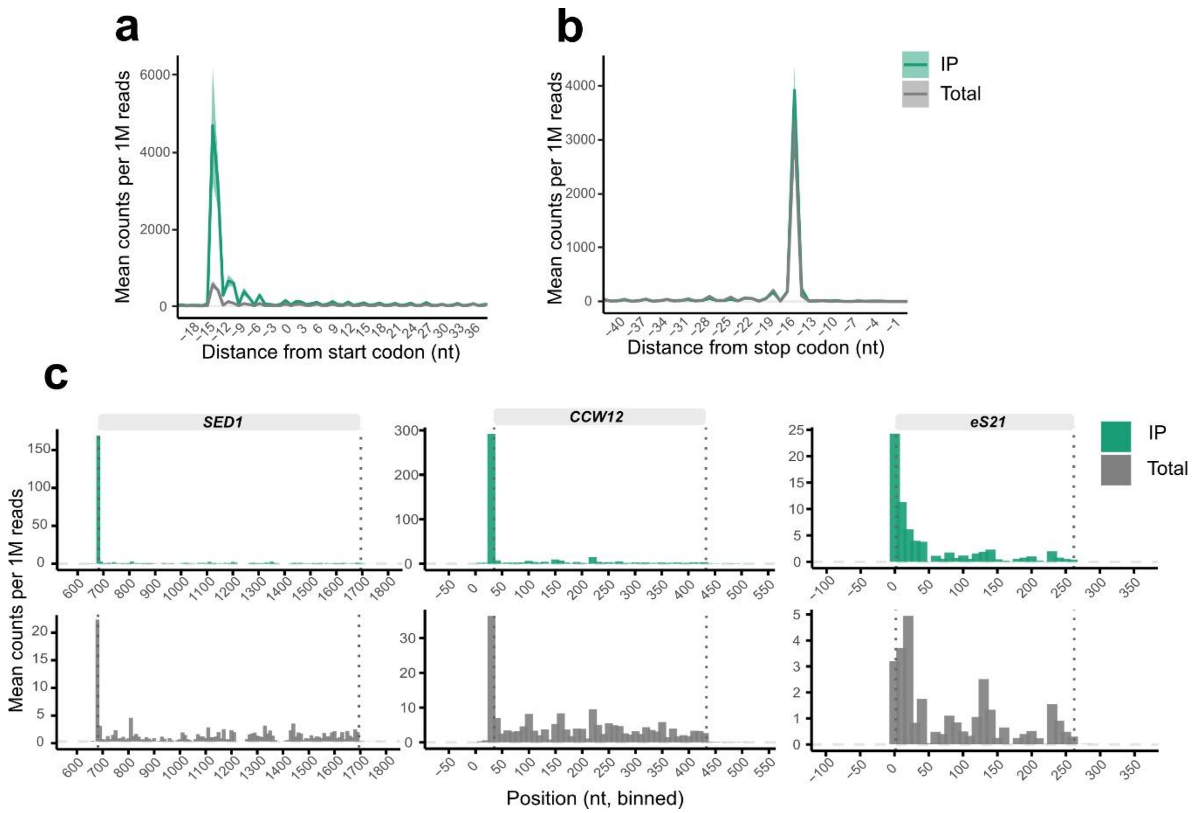


**Supplementary Figure 9: (a, b)** Metagene analysis of ribosome footprints around start codons for Total (grey) and IP (green) samples at 28 nt (a) and 30 nt (b). Read density (counts per million) is plotted as a function of nucleotide position. **(c)** Distribution of 28-nt RPFs on the SED1, CCW12, and eS21 transcripts from purified uL16$^{mut}$ IP and Total ribosome libraries. Data are P-site adjusted. Dotted lines indicate annotated translation start and stop sites.

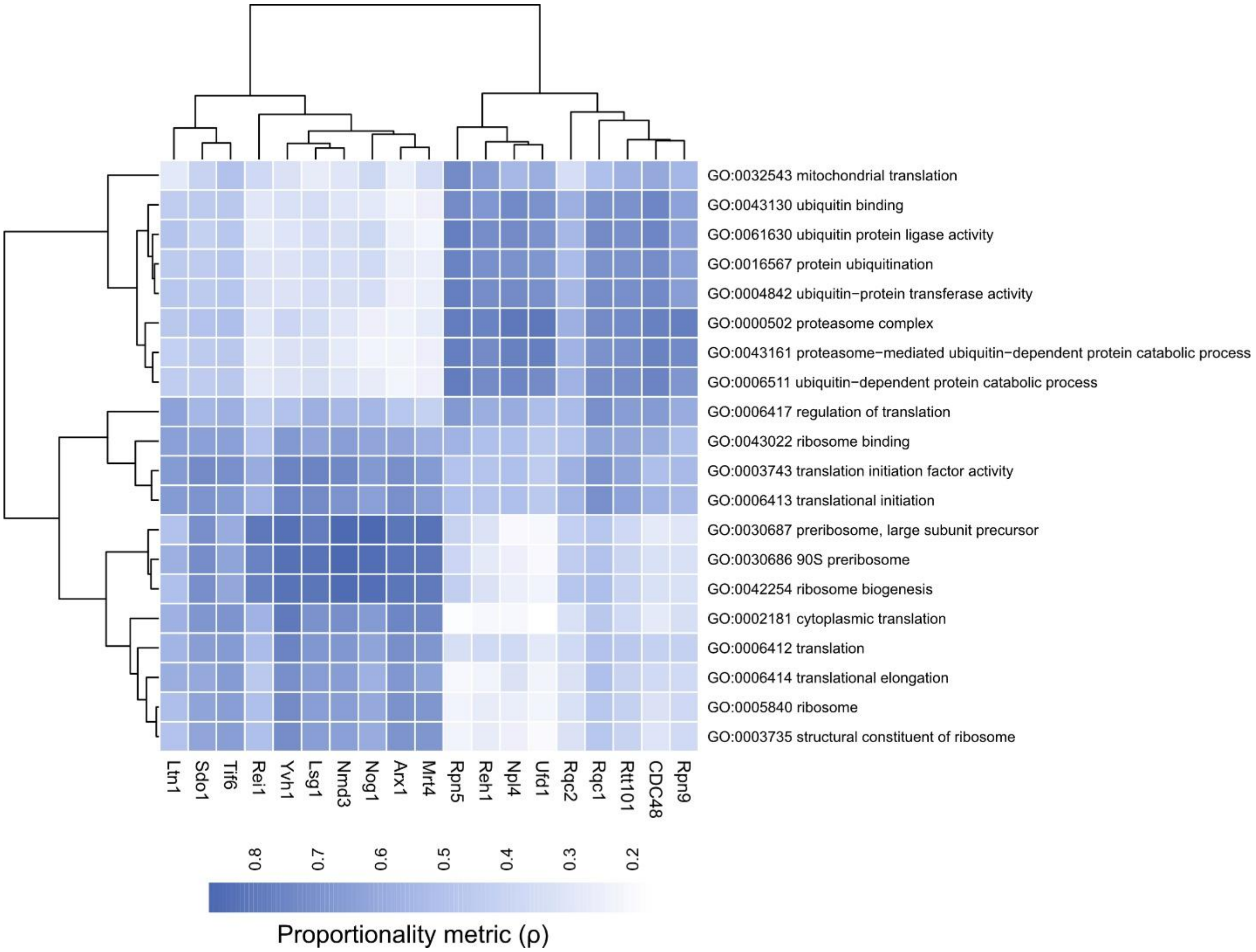


**Supplementary Figure 10: Reh1 co-expression resembles quality control and proteasomal factors.** Heatmap showing median proportionality metric (ρ) for Reh1, ribosome biogenesis factors, quality control factors, and proteasomal components across selected GO terms.
